# S-phase independent silencing establishment in *Saccharomyces cerevisiae*

**DOI:** 10.1101/2020.05.13.094888

**Authors:** Davis Goodnight, Jasper Rine

## Abstract

The establishment of silent chromatin, a heterochromatin-like structure at *HML* and *HMR* in *Saccharomyces cerevisiae*, depends on progression through S phase of the cell cycle, but the molecular nature of this requirement has remained elusive despite intensive study. Using high-resolution chromatin immunoprecipitation and single-molecule RNA analysis, we found that silencing establishment proceeded via gradual repression of transcription in individual cells over several cell cycles, and that the cell-cycle-regulated step was downstream of Sir protein recruitment. In contrast to prior results, *HML* and *HMR* had identical cell-cycle requirements for silencing establishment, with no apparent contribution from a tRNA gene adjacent to *HMR*. We identified the cause of the S-phase requirement for silencing establishment: removal of transcription-favoring histone modifications deposited by Dot1, Sas2, and Rtt109. These results revealed that silencing establishment was absolutely dependent on the cell-cycle-regulated interplay between euchromatic and heterochromatic histone modifications.

## INTRODUCTION

Inheritance of gene expression state often accompanies the inheritance of genetic content during cell division. Indeed, the eukaryotic replication fork plays host to the enzymes needed to replicate DNA as well as intricate machinery that reassembles chromatin in the wake of replication. However, during development, cell division is also coupled to the rewiring of gene expression patterns that lead to the generation of new cell types. An understanding of chromatin and epigenetics requires an understanding of the mechanisms by which cells can both faithfully transmit chromatin state through cell division, as well as subvert that inheritance in order to establish new cell types. The silent chromatin controlling mating-type identity in *Saccharomyces cerevisiae* offers a tractable context for exploring how cell-cycle-regulated chromatin dynamics lead to the establishment of new expression states.

The maintenance of the correct mating type in *Saccharomyces* relies on both the expression of the **a** or α mating-type genes at the *MAT* locus and the heterochromatin-mediated silencing of copies of those same genes at *HML* and *HMR* (Herskowitz, 1989). Silencing is dependent on the <u>S</u>ilent <u>I</u>nformation <u>R</u>egulator genes, *SIR1-4*, whose study has led to an understanding of how silencing is achieved (Gartenberg and Smith, 2016; Rine and Herskowitz, 1987). *HML* and *HMR* are flanked by DNA sequences termed silencers, which recruit the DNA-binding proteins Rap1, Abf1, and ORC. These in turn recruit the Sir proteins via protein-protein interactions. Sir protein recruitment to silencers is followed by the spread of Sir proteins across the multi-kilobase loci by iterative cycles of deacetylation of the tails of histones H3 and H4 by Sir2 and binding of Sir3 and Sir4 to those deacetylated histone tails(Hecht et al., 1995; Hoppe et al., 2002; Rusché et al., 2002).

Despite decades of work, a longstanding puzzle remains at the heart of the mechanism of silencing: cells must pass through S phase to establish silencing, but the identity of the elusive cell-cycle-dependent component is unknown (reviewed in Young and Kirchmaier, 2012). Cells with a temperature-sensitive *sir3-8* allele arrested in G1 cannot repress *HMRa1* when switched from the non-permissive temperature to the permissive temperature, but can when allowed to progress through the cell cycle (Miller and Nasmyth, 1984). DNA replication *per se* is not required for silencing establishment, as excised DNA circles bearing *HMR* but no origin of replication, can be silenced if allowed to pass through S phase (Kirchmaier and Rine, 2001; Li et al., 2001). Thus, some feature of S-phase, but not DNA replication itself, is crucial for silencing establishment.

Interestingly, low-resolution chromatin immunoprecipitation (ChIP) studies showed that Sir protein recruitment to *HMR* can occur with or without cell-cycle progression, suggesting that Sir protein binding and silencing are not inextricably linked (Kirchmaier and Rine, 2006). If Sir proteins can bind to a locus but not silence it, then other molecular changes must be required to create silencing-competent chromatin. In cycling cells undergoing silencing establishment, removal of histone modifications associated with active transcription occurs over several cell cycles (Katan-Khaykovich and Struhl, 2005). Furthermore, deletion genes encoding enzymes that deposit euchromatic histone marks modulate the speed of silencing establishment in cycling cells (Katan-Khaykovich and Struhl, 2005; Osborne et al., 2009). It is unknown whether the removal of euchromatic marks is related to the S-phase requirement for silencing establishment.

To better understand how chromatin transitions from the active to repressed state are choreographed, we developed an estradiol-regulated Sir3 fusion protein which, combined with high-resolution ChIP and RNA measurements, allowed precise experimental analysis of silencing establishment with single-cell resolution. We characterized the molecular changes that occur during silencing establishment and identified the genetic drivers of the S-phase requirement for silencing establishment.

## RESULTS

### S phase as a critical window for silencing establishment

Previous studies of silencing establishment have used a variety of strategies to controllably induce silencing establishment, each with their own strengths and weaknesses (see, e.g., Miller and Nasmyth, 1984; Kirchmaier and Rine, 2001; Li, Cheng and Gartenberg, 2001; Lazarus and Holmes, 2011). We sought a new tool to induce silencing that would allow preservation of the structure of the silencers at *HML* and *HMR* and minimally perturb cell physiology upon induction. To do this, we fused the coding sequence of the estrogen binding domain (*EBD*) from the mammalian estrogen receptor α to *SIR3*, making *SIR3*’s function estradiol-dependent (**Figure 1A**; Lindstrom and Gottschling, 2009; Picard, 1994). Estradiol addition frees the EBD from sequestration by Hsp90, and hence the induction is rapid, because it does not require new transcription or translation (McIsaac et al., 2011). *SIR3-EBD* strains grown without estradiol failed to repress *HMR*, mimicking the *sir3*Δ phenotype, while those grown with estradiol repressed *HMR* to a similar degree as wild-type *SIR3* strains (**Figure 1B**).

**Figure 1:**
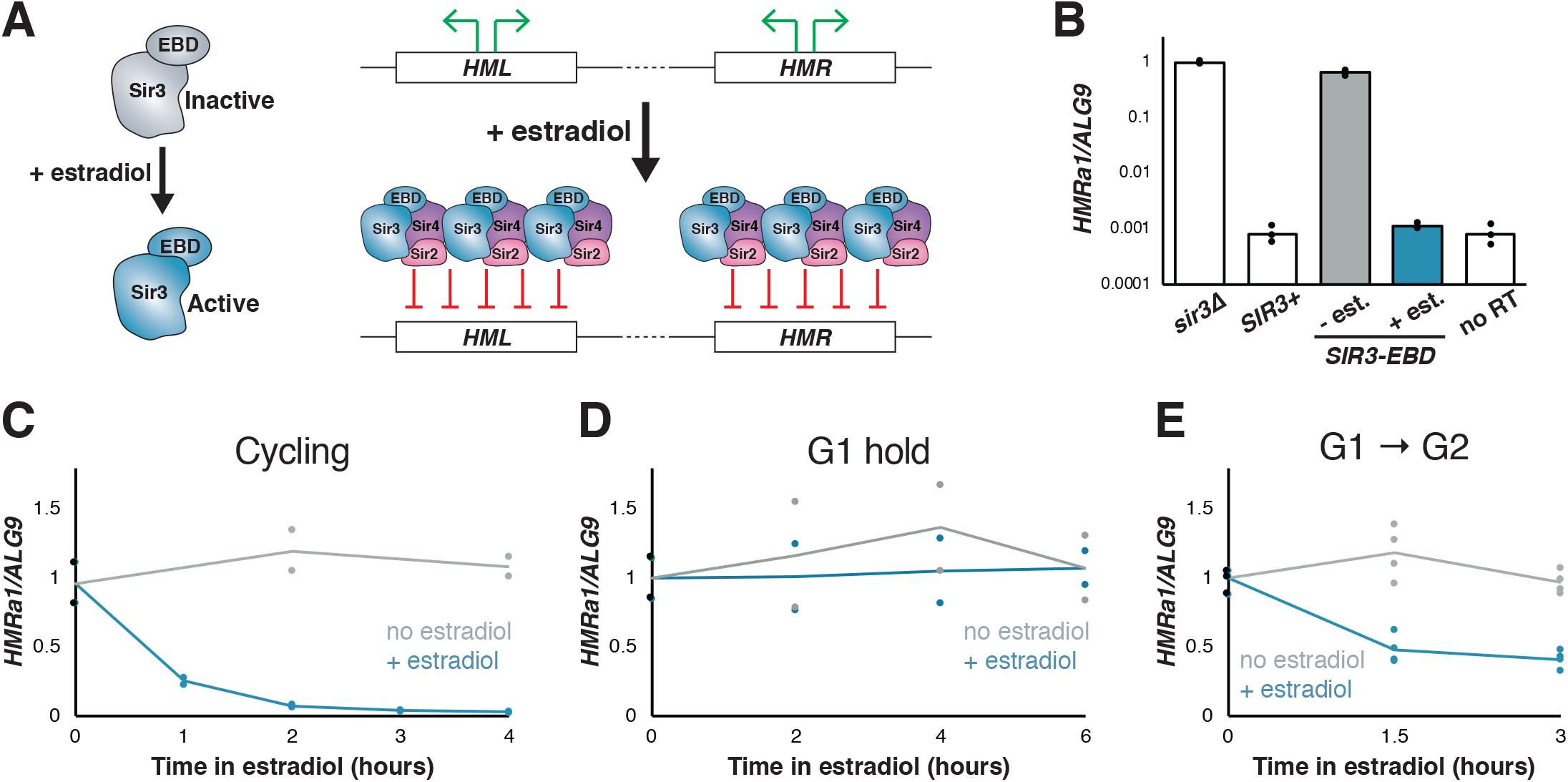
Silencing establishment using *SIR3-EBD* required S-phase progression. **(A)** Schematic for *SIR3-EBD* activation. When estradiol is absent, *SIR3-EBD* is kept inactive and *HML* and *HMR* are expressed. Upon addition of estradiol, *SIR-EBD* is activated and *HML* and *HMR* are repressed. **(B)** RT-qPCR of mRNA from *sir3Δ* (JRY12168), *SIR3+* (JRY12171), and *SIR3-EBD* (JRY12170) cells grown with ethanol (solvent control) or estradiol (N=3 for each condition). The *HMRa1*^no RT^/*ALG9* value for *SIR3+* cells established that *SIR3+* cells and *SIR3-EBD* cells grown with estradiol silenced *HMRa1* to essentially the limit of detection. **(C)** *SIR3-EBD* cultures (JRY12169, JRY12170) were grown to mid-log phase, then split and grown in medium with either estradiol or ethanol added. Silencing was monitored by qPCR in a time course after estradiol addition. t=0 represents the point of estradiol addition for this and subsequent experiments. **(D)** *SIR3-EBD* cultures (JRY12169, JRY12170) were arrested in G1 with α factor, then split, with either ethanol or estradiol added. The arrest was maintained for 6 hours, and silencing was assayed by RT-qPCR throughout. **(E)** *SIR3-EBD* cultures (JRY12169, JRY12170; 2 replicates of each genotype) were arrested in G1 with α factor, then split and released to G2/M by addition of protease and nocodazole in the presence of either ethanol or estradiol. In this and all subsequent figures, dots represent biological replicates, and the bars/lines represent the averages of biological replicates.

To test whether the *SIR3-EBD* allele retained the requirement for cell-cycle progression to repress *HMR*, estradiol was added to cells that were either cycling or arrested in G1 by α factor. In cycling cells, silencing establishment of *HMR* occurred gradually over several hours (**Figure 1C**). However, in cells arrested in G1, estradiol led to no measurable repression of *HMR*, even after many hours (**Figure 1D**). Thus, silencing of *HMR* could not occur without progression through the cell cycle, in agreement with prior results using other conditional alleles.

Prior work indicated that S phase is a critical window during which cells may undergo partial silencing establishment (Kirchmaier and Rine, 2001; Lau et al., 2002; Miller and Nasmyth, 1984). Consistent with this, when we arrested cells in G1, then induced *SIR3-EBD* and allowed them to proceed through S phase and re-arrested them at G2/M, *HMR* was repressed ∼60% from its starting levels (**Figure 1E**). Crucially, the extent of this partial repression was stable over many hours in these G2/ M-arrested cells. Thus, a repression-permissive window or event occurred between G1 and the beginning of mitosis that allowed partial silencing establishment, and further repression was not possible while arrested at G2/M. Indeed, after 3 hours in estradiol, cycling cells were repressed >20-fold from their starting value, compared to only ∼3-fold for cells arrested after a single S phase (compare **Figures 1C** and **1E**). This requirement for multiple cell cycles to occur before full gene repression was achieved was consistent with prior studies of silencing establishment both in cell populations and at the single-cell level (Katan-Khaykovich and Struhl, 2005; Osborne et al., 2009).

Having established the validity of the *SIR3-EBD* fusion as a tool for studying silencing, we revisited two mutants that have been reported to bypass cell-cycle requirements for silencing establishment. In one study, depletion of the cohesin subunit Mcd1/Scc1 allowed for increased silencing in G2/M-stalled cells (Lau, Blitzblau & Bell 2002). In another study, deletion of a tRNA gene adjacent to *HMR*, termed *tT(AGU)C*, which is known to bind cohesin, was found to allow partial silencing establishment at *HMR* without S-phase progression (Lazarus and Holmes 2011). Using *SIR3-EBD* in combination with an auxin-inducible degron (AID)-tagged Mcd1, we found no effect of depleting cohesin or deleting the tRNA gene in regulating silencing establishment (**Figure 1—figure supplement 1**). Thus, at least for strains using *SIR3-EBD*, the genetic basis for the cell-cycle requirement for silencing establishment at *HMR* was unknown prior to the work described below. Possible explanations for the discrepancies between our results and earlier reports are discussed below.

Our finding that *tT(AGU)C* did not regulate silencing establishment led us to reconsider the broader claim that *HMR* is distinct from *HML* in its requirement for S phase for silencing establishment (Lazarus and Holmes, 2011; Ren et al., 2010). Earlier silencing establishment assays at *HML* were complicated by the strong silencing-independent repression of *HMLα1* and *HMLα2* by the **a**1/α2 repressor (Herskowitz, 1989; Siliciano and Tatchell, 1986). To avoid this limitation, we constructed an allele of *HML* with nonsense mutations in both *α1* and *α2*, so that the α1 and α2 proteins were never made, even when *HML* was de-repressed. This modification also allowed us to use α factor to arrest cells while studying silencing establishment at *HML*, which was not possible before. We introduced additional single nucleotide polymorphisms into the regions of *HML* that are homologous to *HMR*, to allow unambiguous assignment of high-throughput sequencing reads to one of the two loci (see below). We refer to the mutant locus as *HML\** and the mutant alleles as *hmlα1\** and hml*α2\** hereafter.

When cells with *HML\** and *SIR3-EBD*, were arrested in G1 and then treated with estradiol, they were unable to silence *hmlα1\** or *hmlα2\** while kept in G1 (**Figure 1—figure supplement 2A, 2B**). Interestingly, expression of *hmlα1\** and hml*α2\** increased markedly over the course of the α-factor arrest. This α-factor-dependent hyper-activation was observed even in *sir3*Δ cells in which no silencing occurs (**Figure 1—figure supplement 2E, 2F**). We identified two previously-unreported binding sites for Ste12, the transcription factor activated by mating pheromone, in the bidirectional *α1*/*α2* promoter, which explains the increased expression when cells are exposed to α factor (**Figure 1—figure supplement 2G**; Dolan, Kirkman and Fields, 1989). Both *hmlα1\** and *hmlα2\** decreased in expression following release from G1 to G2/M (**Figure 1—figure supplement 2C, 2D**), suggesting that S phase is required for partial silencing establishment at *HML*. Notably, the fold change in expression that followed a single passage through S phase was not identical among *HMRa1*, *hmlα1\**, and *hmlα2\**. Thus, some S-phase-dependent process is important for silencing both *HML* and *HMR*, but the effects of that process vary in magnitude between these two loci.

### Silencing establishment occurred through a partially repressive intermediate

We were interested in the partial silencing observed in cells that transited through a single S phase after *SIR3-EBD* induction, in which transcription of *HMRa1* was down ∼60% (see **Figure 1E**). This appearance of a stable intermediate level of silencing could reflect either of two distinct phenomena at the single-cell level (**Figure 2A**). One possibility is that cells have a ∼60% chance of establishing stable heterochromatin during the first S phase after *SIR3-EBD* induction and a ∼40% chance of failing to do so. This would resonate with the behavior of certain mutants, e.g. *sir1*Δ, wherein silent loci can exist in one of two epigenetic states: stably repressed or stably de-repressed, with rare transitions between the two (Pillus and Rine, 1989; Xu et al., 2006). Alternatively, every cell might reach a partially repressive chromatin state at *HMR* during the first S phase during silencing establishment.

**Figure 2:**
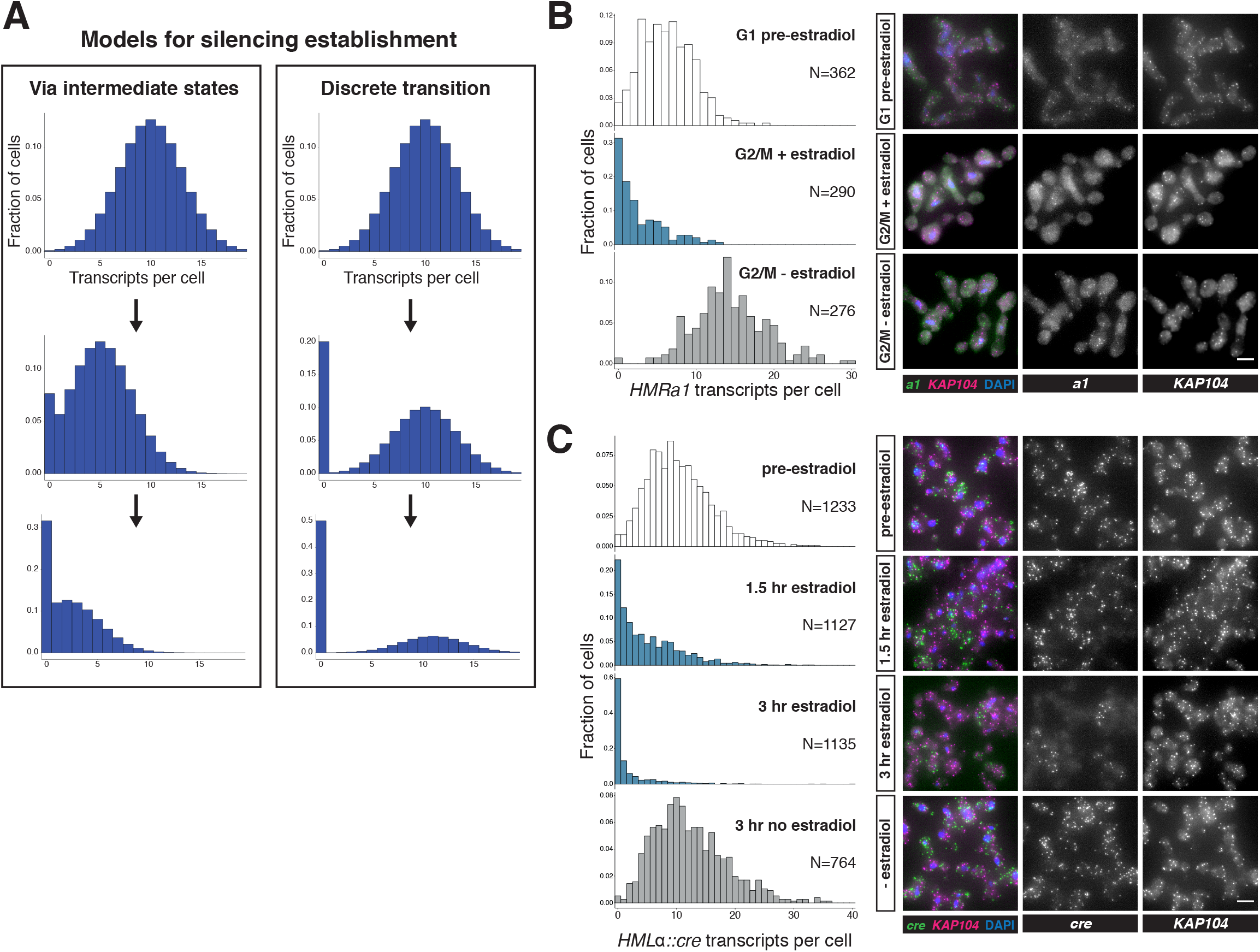
Silencing establishment proceeded via gradual repression in individual cells. **(A)** Potential models for silencing establishment. Before silencing establishment (top), mRNA transcripts are present as a distribution around a mean. If silencing establishment occurred via intermediate states (left), the mean number of transcripts per cell would decrease over time, with complete silencing, i.e., zero transcripts per cell, occurring as the probability distribution shifted toward the y axis. If silencing establishment occurred via discrete transitions (right), an increasing fraction of cells would have zero transcripts over time, but the distribution of cells with >0 transcripts would retain the same shape. **(B)** smRNA-FISH for *HMRa1* during silencing establishment after 1 S phase. A *SIR3-EBD* culture (JRY11762) was arrested in G1 with α factor (“G1 pre-estradiol”), then split and released to G2/M by addition of protease and nocodazole in the presence of either estradiol or ethanol. Samples were collected 2 hours after estradiol addition. **(C)** smRNA-FISH for *HMLα∷cre* during silencing establishment in cycling cells. A *SIR3-EBD* strain bearing the *HMLα∷cre* reporter (JRY12514) was grown to mid-log phase (“pre-estradiol”), then the culture was split in two, with one sub-culture receiving estradiol and the other receiving ethanol. Samples were collected for smRNA-FISH at t=1.5 hours and t=3 hours after estradiol addition. For both (B) and (C), the images displayed are representative maximum-intensity Z-projections. The data shown in (B) and (C) each represent one of two replicate experiments, for which the other replicate is shown in Figure 2—figure supplements 1 and 2, respectively. Scale bars = 5 *μ*m.

To distinguish between these possibilities, we used single-molecule RNA fluorescence in-situ hybridization (smRNA-FISH) to quantify the expression of *HMRa1*during the establishment process. If silencing establishment proceeded via individual cells transitioning between the discrete “ON” and “OFF” states during S phase, we would expect an accumulation of cells with zero transcripts during the establishment of silencing, with no change in the average number of transcripts in those cells still expressing *HMRa1*. However, if silencing establishment proceeded via partially repressive intermediates in individual cells, we would expect a shift downward in the mean number of transcripts per cell (**Figure 2A**). In both cases, cells with zero transcripts would accumulate over time.

As expected, *SIR3-EBD* cells arrested in G1 without estradiol had similar numbers of *HMRa1* transcripts as *sir3*Δ cells (**Figure 2—figure supplement 1A**). When we added estradiol and allowed the cells to go through S phase to G2/M, the decrease in transcript number in the population of cells analyzed closely mirrored the results we obtained using RT-qPCR, confirming that our single-molecule analysis was consistent with bulk measurements (**Figure 2—figure supplement 1B**, compare to **Figure 1E**). This decrease occurred via a reduction in the average number of transcripts per cell, and not simply by an increase in the number of cells with zero transcripts (**Figure 2B**, **Figure 2—figure supplement 1C, 1D**). Thus, individual cells undergoing silencing establishment at *HMR* formed partially repressive heterochromatin after a single S phase.

To determine whether the stepwise repression seen at *HMR* was a general feature of silencing establishment or if it was particular to *HMR* and/or the cell-cycle conditions tested, we performed an analogous experiment using the *HMLα∷cre* allele that has been previously characterized by smRNA-FISH (Dodson and Rine, 2015). In this experiment, we analyzed the number of *cre* transcripts per cell over time following induction of *SIR3-EBD*, without any cell-cycle perturbations. Silencing establishment at *HML* also occurred via partially repressive intermediate states (**Figure 2C**, **Figure 2—figure supplement 2A, 2B**). The degree of repression observed by smRNA-FISH was quantitatively similar to the measurement of the same gene by RT-qPCR (**Figure 2—figure supplement 2C, 2D**). Together, these results suggested that silencing establishment proceeds via the gradual repression of genes by the Sir proteins, and that there are specific windows of the cell cycle during which transcriptional tune-down can occur.

### Extensive Sir protein binding could occur without gene repression

The gradual silencing establishment described above might be achieved via increased Sir protein recruitment to *HML* and *HMR* during each passage through a specific cell-cycle window. Alternatively, Sir protein recruitment might be independent of the cell cycle, in which case passage through the cell cycle would favor repression via a step downstream of Sir protein recruitment. To test whether Sir protein recruitment was limited in the cell cycle, we performed ChIP-seq on myc-tagged Sir4 during silencing establishment. The tagged Sir4-myc is functional for silencing (**Figure 1B**), and its localization at *HML* and *HMR* is indistinguishable from Sir2-myc and Sir3-myc in wild-type cells (Thurtle and Rine, 2014).

As expected, ChIP-seq on cells with wild-type *SIR3* revealed strong binding of Sir4-myc throughout *HML\** and *HMR* (**Figure 3A** & **3B**). *SIR3-EBD* cells grown with estradiol gave profiles that were indistinguishable in both the strength of binding and the location of binding. Surprisingly, in *sir3*Δ cells and *SIR3-EBD* cells grown without estradiol, Sir4-myc binding at *HML\** and *HMR,* though severely reduced, was still evident (**Figure 3A**). This weak binding was observed in multiple replicates with different crosslinking times and was not observed in cells with untagged Sir4 (**Figure 3—figure supplement 1A**). At neither *HML\** nor *HMR* was there evidence for preferable Sir4-myc binding at the E or I silencers in absence of *SIR3*. This distribution was in marked contrast with the expectation of the earlier model of nucleation and spreading of heterochromatin, as discussed below.

**Figure 3:**
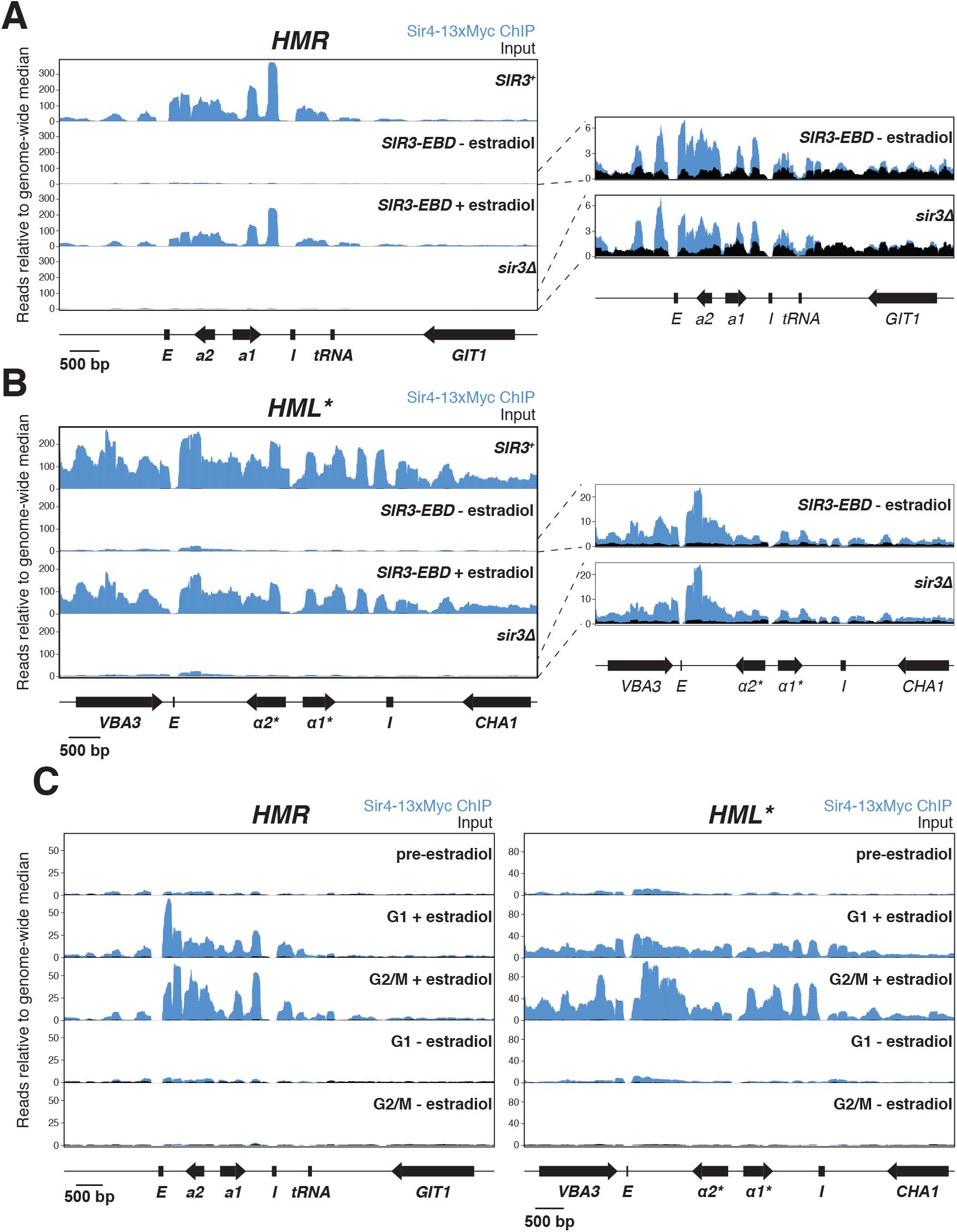
Sir protein binding and silencing were separable phenomena. All panels show Sir4-13xMyc ChIP-seq signal in blue and input in black. Read counts were normalized to the non-heterochromatin genome-wide median. IP and input values are plotted on the same scale. **(A)** Left, ChIP-seq for Sir4-13xMyc at *HMR* in strains with *SIR3+* (JRY12172), *sir3Δ* (JRY12168), and *SIR3-EBD* (JRY12170) grown with or without estradiol and fixed for 60 minutes in formaldehyde. Right, same data as the left panel for *sir3Δ* and *SIR3-EBD* without estradiol, enlarged to show IP levels above input. **(B)** Same as (A), but showing data from *HML\**. **(C)** ChIP-seq for Sir4-13xMyc during silencing establishment at *HMR* (left) and *HML\** (right). Cultures of *SIR3-EBD* cells (JRY12169) were arrested in G1 with α factor (“pre-estradiol”), then split four ways. Two sub-cultures were maintained in G1 in medium with estradiol or ethanol (“G1 + estradiol” and “G1 - estradiol”). The other two sub-cultures were released to G2/M by addition of protease and nocodazole; and received either estradiol or ethanol (“G2/M + estradiol” and “G2/M − estradiol”). After 3 hours in medium with estradiol or ethanol, cultures were fixed in formaldehyde for 15 minutes and collected for ChIP-seq. Data shown represent one of two replicates, with the other shown in Figure 3—figure supplement 1C.

ChIP-seq data from *SIR3-EBD* cells arrested in G1 without estradiol revealed the same weak enrichment of Sir4-Myc at *HML\** and *HMR* that we observed in cycling cells (**Figure 3C** & **Figure 3—figure supplement 1C**). However, upon addition of estradiol in cells kept in G1, we saw a strong increase in Sir4-myc binding across the loci (**Figure 3C** & **Figure 3—figure supplement 1C**). The increase in Sir4-myc binding to *HMR* was not associated with any change in expression of *HMRa1*, which remained completely de-repressed (**Figure 3—figure supplement 1B**). Hence, Sir protein binding across *HML\** and *HMR* was not sufficient to lead to gene silencing. When cells were allowed to pass from G1 to G2/M, the resulting partial silencing was correlated with an increase in Sir4-myc binding at *HML\** and *HMR* (**Figure 3C** & **Figure 3—figure supplement 1C**). Thus, Sir proteins binding throughout *HML\** and *HMR* in absence of cell-cycle progression achieved no repression, and some S-phase-dependent process promoted further binding and partial repression. Together, these data revealed the existence of cell-cycle-regulated steps beyond Sir binding required to bring about silencing.

### Removal of H3K79 methylation was a critical cell-cycle-regulated step in silencing establishment

Given that silencing at *HML* and *HMR* was established only during a discrete window of the cell cycle, the key issue was to identify what molecular event(s) occurred during this window and why it/they were limited in the cell cycle. A mutant that could establish silencing while arrested in G1 would potentially identify that molecular event.

The histone methyltransferase Dot1 has several characteristics that suggest it might act as an antagonist of silencing establishment. Dot1 methylates histone H3 on lysine 79 (H3K79), which interferes with Sir3 binding to nucleosomes (Altaf et al., 2007; Armache et al., 2011; Van Leeuwen et al., 2002; Yang et al., 2008). Dot1 is unique among yeast histone methyltransferases in lacking a counteracting demethylase that removes H3K79 methylation. Thus, removal of H3K79 methylation can be achieved only through turnover of the histones that bear it, such as occurs during S phase, when new histones are incorporated that lack H3K79 methylation (De Vos et al., 2011). Indeed, *dot1*Δ *SIR3-EBD* cells arrested in G1 robustly repressed *HMRa1*, *hmlα1\**, and *hmlα2\** upon addition of estradiol (**Figure 4A**, **Figure 4—figure supplement 1A, 1B**), suggesting that removal of H3K79me from *HML* and *HMR* was one crucial S-phase-specific step during silencing establishment.

**Figure 4:**
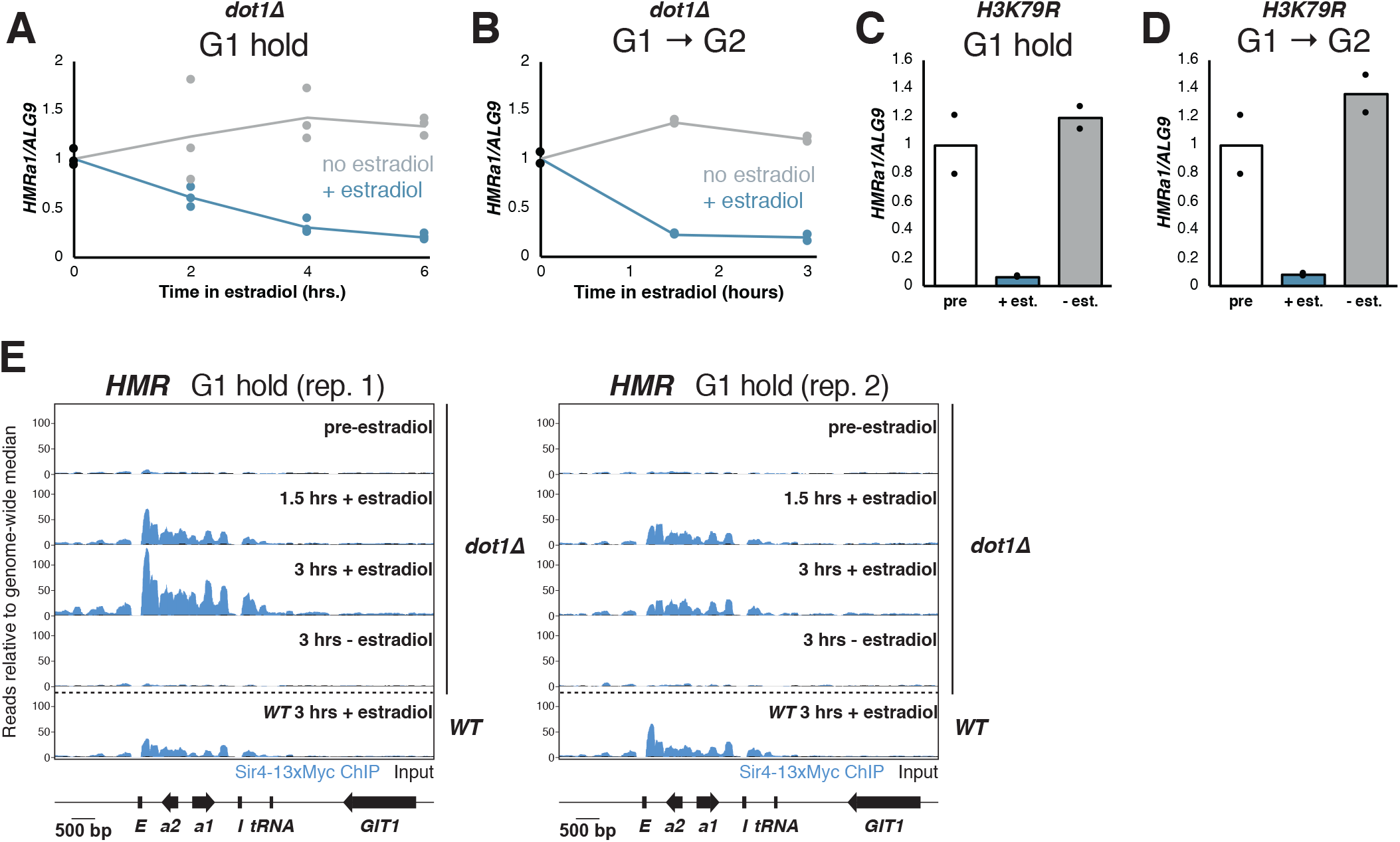
Cells without H3K79 methylation established silencing without cell-cycle progression. **(A)** Cultures of *dot1Δ* cells (JRY12443, JRY12445) were arrested in G1 with α factor, then split, with half receiving ethanol and the other half receiving estradiol. Silencing was monitored by RT-qPCR over time after estradiol addition. **(B)** *dot1Δ* mutants were arrested in G1 with α factor, then released to G2/M by addition of protease and nocodazole, and either ethanol or estradiol. Silencing was monitored by RT-qPCR over time after estradiol addition. **(C)** Cultures of cells in which lysine 79 was mutated to arginine in both *HHT1* and *HHT2* in two isogenic strains (*H3K79R*; JRY12851, JRY12852) were arrested in G1 with α factor, then split, with one sub-culture receiving ethanol and the other receiving estradiol. Silencing was assayed by RT-qPCR after 6 hours in ethanol. **(D)** *H3K79R* cells were arrested in G1 with α factor, then released to nocodazole with protease and nocodazole and either estradiol or ethanol. Silencing was assayed by RT-qPCR after 3 hours in estradiol. The pre-estradiol sample for this experiment was the same culture used in (C). **(E)** Cultures of *dot1Δ* cells (JRY12443, JRY12444) were arrested in G1 with α factor (“pre-estradiol”), then split, with half the culture receiving ethanol, and the other half receiving estradiol. After 1.5 hours and after 3 hours, samples were fixed for 15 minutes in formaldehyde and collected for ChIP. Sir4-13xMyc ChIP-seq signal is in blue and input in black, each normalized to the non-heterochromatin genome-wide median and plotted on the same scale. Also displayed are two replicates of wild-type G1 cells after 3 hours in estradiol from Figures 3C and Figure 3—figure supplement 1C.

To test whether the *dot1*Δ phenotype was due specifically to methylation at H3K79, both copies of histone H3 were mutated to encode arginine at position 79 (*H3K79R*), a mimic for the non-methylated state. This mutant also allowed for robust silencing establishment in G1, in fact, to a stronger degree than *dot1*Δ (**Figure 4C**). Strains with H3K79 mutated to leucine (*H3K79L*) or methionine (*H3K79M*) failed to establish silencing even after passage through S phase (**Figure 4—figure supplement 1C, 1D**), confirming the importance of the positive charge on H3K79 in silencing. Notably, even though G1-arrested *H3K79R* cells could strongly repress *HMRa1* (∼15-fold), this was still incomplete relative to fully silenced cells, which repressed *HMRa1* >1000-fold (see **Figure 1B**). Thus, either increased time or cell cycle progression promoted silencing establishment even in absence of H3K79 methylation.

In addition to promoting S-phase-independent silencing establishment, *dot1*Δ and *H3K79R* cells that passed from G1 to G2/ M also repressed *HMRa1* more robustly than did wild-type cells transiting the same cell-cycle window (**Figure 4B & Figure 4D,** compare to **Figure 1E**). S phase passage markedly increased the speed of silencing establishment in *dot1*Δ cells, though the ultimate degree of repression was similar whether cells passed through S phase or stayed in G1 (compare **Figures 4A & 4B**). Thus, S phase still promotes silencing establishment in cells lacking H3K79 methylation.

To understand how H3K79 methylation prevented silencing establishment, we performed ChIP-seq for Sir4-myc in *dot1*Δ cells undergoing silencing establishment in the absence of cell-cycle progression. G1-arrested *dot1*Δ cells already displayed clear partial silencing establishment after 1.5 hours in estradiol (**Figure 4—figure supplement 1E**). However, the level of Sir4-myc recruitment to *HML\** and *HMR* at this early time point was similar to the recruitment observed in wild-type cells, in which no gene repression had occurred after 3 hours (**Figure 4E** & **Figure 4—figure supplement 1F**). Therefore, even though *dot1*Δ and wild-type cells had indistinguishable levels of Sir binding, that binding gave rise to different transcriptional effects. Thus, removal of removal of H3K79me did not regulate silencing establishment by controlling Sir protein binding.

### SAS2 and RTT109 contributed to limiting silencing establishment to S phase

The crucial role of H3K79 methylation removal in silencing establishment led us to consider other chromatin modifications that might regulate silencing establishment. Two histone acetyltransferases, Sas2 and Rtt109, were especially dsinteresting given the S-phase dynamics of the marks they deposit and their known relevance to silencing. Sas2, the catalytic component of the SAS-I complex, acetylates H4K16 during S phase (Kimura et al., 2002; Meijsing and Ehrenhofer-Murray, 2001; Reiter et al., 2015; Suka et al., 2002). The removal of H4K16 acetylation by Sir2 is the central histone modification associated with silencing (Imai et al., 2000a; Johnson et al., 1990; Landry et al., 2000; Park and Szostak, 1990). Rtt109 acetylates newly-incorporated histone H3 at lysines 9 and 56 during S phase, and this acetylation is largely removed by Hst3 and Hst4 by the time of mitosis (Adkins et al., 2007; Celic et al., 2006; Driscoll et al., 2007; Fillingham et al., 2008; Schneider et al., 2006). Mutations in *SAS2* and *RTT109* have both been shown to have silencing phenotypes (Imai et al., 2000b; Miller et al., 2008).

Interestingly, both *sas2*Δ and *rtt109*Δ mutations led to partial repression of *HMR* upon *SIR3-EBD* induction in cells arrested in G1 (**Figure 5A**). The magnitude of this effect was weaker than in *dot1*Δ cells but highly significant. Cells without *RTT109* grew slowly and were less sensitive to α factor than wild-type cells, so we cannot exclude the possibility that a population of *rtt109*Δ cells passed through S phase during the experiment, contributing to the observed phenotype (**Figure 5—figure supplement 1**). When combined with *dot1*Δ, both *sas2*Δ and *rtt109*Δ led to a further increase in silencing establishment. Thus, *SAS2* and *RTT109* impeded silencing establishment by a different mechanism than *DOT1*. Silencing establishment in cells lacking both *SAS2* and *RTT109* was not significantly different from that of the single mutants. Strikingly, triple mutant *sas2*Δ *rtt109*Δ *dot1*Δ strains established silencing no better than single-mutant *dot1*Δ cells. Interestingly, the G1 phenotypes we observed at *HMR* for *dot1*Δ, *sas2*Δ, and *rtt109*Δ single mutant cells were largely similar at *hmlα1\** (**Figure 5B**). Altogether, these findings demonstrate that *SAS2*, *RTT109*, and *DOT1* inhibit silencing establishment outside of S phase.

**Figure 5:**
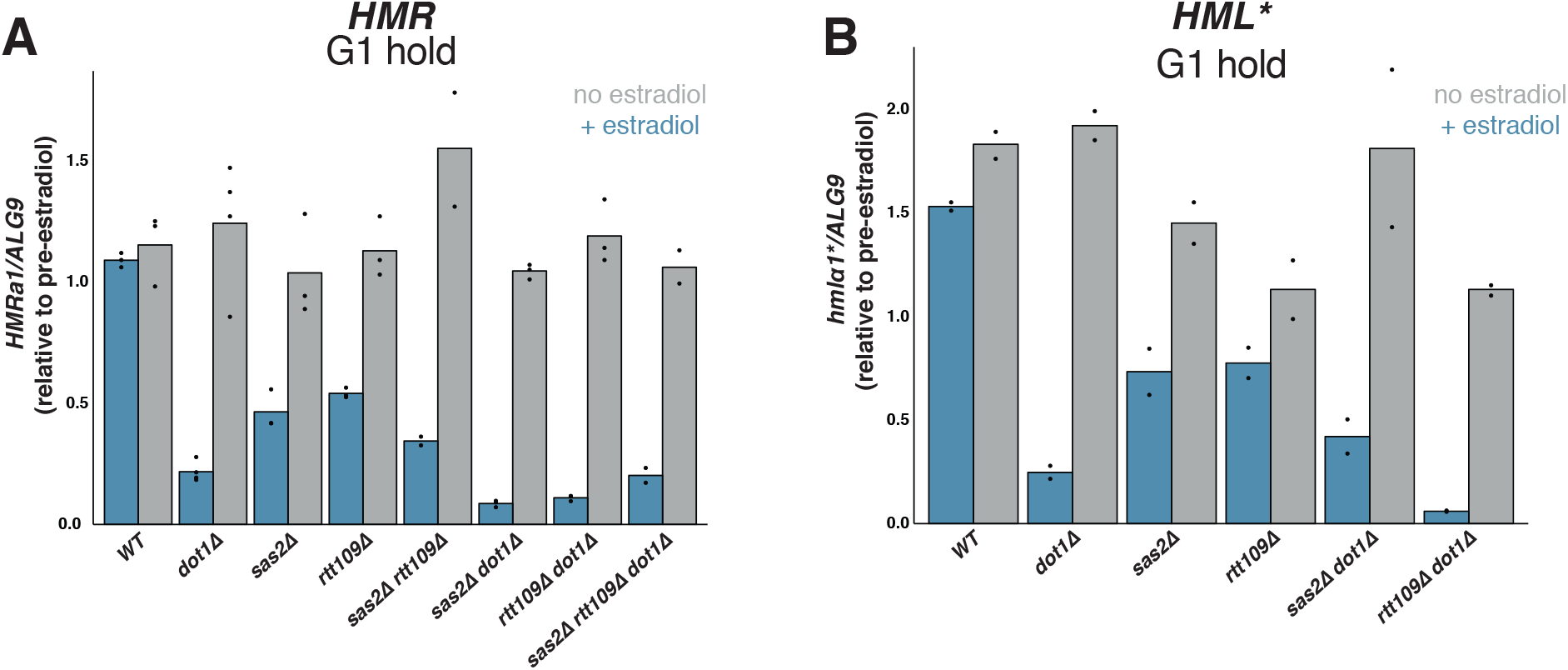
Effects of *SAS2* and *RTT109* on silencing establishment in G1. For all strains, cultures were arrested in G1 with α factor, then split, with one sub-culture receiving estradiol and the other receiving ethanol. Silencing was assayed by RT-qPCR 6 hours after additions. Each sample was normalized to its own pre-estradiol value. The following strains were used. WT: JRY12169 (the parent strain for all mutants); *dot1Δ*: JRY12443, JRY12445; *sas2Δ*: JRY12615, JRY12616; *rtt109Δ*: JRY12689, JRY12690; *sas2Δ rtt109Δ*: JRY12765, JRY12766; *sas2Δ dot1Δ*: JRY12618, JRY12619; *rtt109Δ dot1Δ*: JRY12691, JRY12692; *sas2Δ rtt109Δ dot1Δ*: JRY12767, JRY12768. **(A)** Silencing establishment of *HMRa1* by RT-qPCR. The level of repression observed in each mutant was significantly greater than in wild type (Two-tailed T-test; p < 0.005 for each pair-wise comparison). The level of repression observed in *sas2Δ dot1Δ* and *rtt109Δ dot1Δ* double mutants was significantly greater than in the *dot1Δ* single mutant (p < 0.01 for each pair-wise comparison), but there was no significant difference between the values from the *dot1Δ* single mutant and the triple mutant *sas2Δ rtt109Δ dot1Δ* (p = 0.70). **(B)** Silencing establishment of *hmlα1\** by RT-qPCR in a subset of mutant strains. The level of repression for each mutant was significantly greater than in wild type (p < 0.05 for each pair-wise comparison). The level of repression observed in the *rtt109Δ dot1Δ* double mutant was significantly greater than in the *dot1Δ* single mutant (p = 0.028), but there was no significant difference between the values from *sas2Δ dot1Δ* double mutant and the *dot1Δ* single mutant (p = 0.19).

## DISCUSSION

In this study, we identified why silencing establishment requires cell cycle progression. These results highlight the value of studying the dynamics of silencing both in populations of cells and at the single cell level. By monitoring changes in chromatin and changes in expression simultaneously, we documented effects that were elusive at steady state, but critical for a mechanistic understanding of the process. We found that the cell-cycle-dependent removal of euchromatic marks was a major driver of a cell’s ability to establish stable heterochromatin. Interpretation of our results required critical reassessment of some earlier results.

### Silencing establishment occurred by tuning down transcription in individual cells after Sir proteins were bound

The classic model for silencing establishment involves two steps: nucleation of Sir proteins at the silencers, followed by spreading of Sir proteins from silencers via the stepwise deacetylation of nucleosomes by Sir2 and subsequent binding of Sir3 and Sir4 to deacetylated positions of H3 and H4 tails (Hecht et al., 1995; Hoppe et al., 2002; Rusche et al., 2003; Rusché et al., 2002). In the classic model, individual Sir proteins are recruited to the silencers, but the spread across the locus is dependent on all three proteins Sir2/3/4. The binding of a Sir2/3/4 complex to internal nucleosomes at *HML* and *HMR* is thought to drive gene repression at least partly through sterically preventing other proteins from accessing the underlying DNA (Loo and Rine, 1994; Steakley and Rine, 2015). A puzzling observation is that, qualitatively, the nucleation and spread of Sir2, Sir3, and Sir4 to *HML* and *HMR* appears to be cell-cycle-independent, even though the silencing activity of these proteins is clearly dependent on cell cycle progression (Kirchmaier and Rine, 2006).

Surprisingly, we found no condition in which Sir proteins bound only to the silencers. Rather, silencing establishment led to increased Sir4 binding more or less uniformly across the silent loci, beginning from a low-level distributed binding that was present even in the absence of Sir3. Because our ChIP protocol used MNase to fragment chromatin, we observed an under-recovery of DNA not bound by nucleosomes. Thus, we cannot exclude the possibility that our data under-represent the enrichment of Sir4 at the silencers, which are bound by Rap1, Abf1, and ORC, rather than being wrapped around a histone octamer. Even if that were the case, the weak Sir4 binding across *HML* and *HMR* in *sir3*Δ cells suggested that the full Sir2/3/4 complex was not necessary for the distributed binding of Sir4. Rather, Sir3 appeared to stabilize or otherwise enhance Sir4-nucleosome interactions. Further high-resolution ChIP-seq studies will be needed to determine whether Sir2, Sir3, and Sir4, which localize indistinguishably in wild-type cells (Thurtle and Rine, 2014), behave similarly in absence of the full complex, and during silencing establishment.

Upon induction of *SIR3-EBD*, G1-arrested wild-type cells could robustly recruit Sir4 to *HML* and *HMR* without causing any gene repression at these loci. Passage through S phase led to increased Sir binding and partial silencing establishment. However, in G1-arrested *dot1*Δ cells, in which Sir4 binding patterns were indistinguishable from G1-arrested wild-type cells, induction of *SIR3-EBD* caused partial silencing establishment. Together, these observations indicate that a key regulated step in building heterochromatin occurred after the major silencing factors were already present at the locus. Two interpretations were compatible with our data. First, the non-repressive Sir4 binding observed in G1 and the repressive Sir4 binding observed in G2/M could differ in some parameter that is not apparent in crosslinking ChIP experiments, such as differences in the on and off rates for Sir4 binding to nucleosomes. Second, Sir binding could be unable to drive transcriptional changes until competing euchromatic marks on chromatin are relieved. Consistent with the latter interpretation, a prior study of telomeric silencing found that Sir protein binding was detectable at both repressed and de-repressed telomeres, but euchromatic marks, including H3K79me, were found only at de-repressed telomeres (Kitada et al., 2012).

Our smRNA-FISH results showed that silencing establishment proceeds via the gradual tune-down of transcription in individual cells, and that this tune-down occurs over multiple cell cycles. This result conflicts with a prior study of silencing establishment in single cells using a fluorescent reporter at *HML*. That study concluded that silencing establishment proceeded via discrete transitions from the “ON” to the “OFF” state (Xu et al., 2006). However, because that study relied on qualitative assessment of fluorescence intensity in individual cells, it may not have been possible to ascertain intermediate states. Indeed, our data illustrate an inherent limitation of qualitative measurements of single-cell parameters: in the smRNA-FISH images in **Figure 2**, a striking feature is the dichotomy between cells with no transcripts and those with some transcripts. That observation might lead to the conclusion that silencing establishment is caused by the complete shutdown of transcription stochastically in some cells. However, as illustrated by **Figure 2A**, that dichotomy is expected from both an “all-or-nothing” model and a “gradual transition” model. It is only through the quantitative analysis that we could see the gradual decrease in transcription in individual cells.

Whether silencing acts through steric occlusion or through a more specific inhibition of some component necessary for transcription, it is difficult to explain how any intermediate in the assembly of a static heterochromatin structure could drive partial repression. The simplest explanation for how partially repressive chromatin could form would be that silencing machinery and transcriptional machinery both come on and off the chromatin, and the establishment of silencing involves a change in the relative rates of those two processes. In that case, histone modifications could be crucial in shifting the balance.

### Silencing establishment occurred via similar mechanisms at different loci

The mechanism of repression at the two silent mating type loci, *HML* and *HMR*, is generally assumed to be quite similar, but there are mutations that cause effects only at one of the two loci, and others that cause divergent phenotypes between the two loci (Park and Szostak, 1990; Ehrenhofer-Murray, Rivier and Rine, 1997; Yan and Rine, unpublished). Earlier studies concluded that cell-cycle requirements for silencing establishment differed at *HML* and *HMR* (Ren, Wang and Sternglanz, 2010; Lazarus and Holmes, 2011) In contrast, in both wild-type cells and the mutant conditions we tested, both loci behaved similarly. A major innovation that distinguished our studies from the prior studies was our use of a mutant *HML* that allowed unambiguous study of its expression by ensuring that α1 and α2 proteins would not be made. This strategy removed the strong repressive effect that the **a**1/α2 repressor has on transcription from the *HML* promoter, which was a confounding influence in earlier experimental designs that could have led to apparent cell-cycle independent silencing of *HML*.

This fundamental similarity between *HML* and *HMR* in silencing establishment was further evidenced by the lack of an effect of the tRNA gene adjacent to *HMR,* or the tRNA gene’s binding partner, cohesin, loss of either of which were reported to allow early silencing establishment in previous studies (Lau et al., 2002; Lazarus and Holmes, 2011). The reason behind the difference between our results and those of the previous studies is not clear. We did observe subtle silencing-independent fluctuations in *HMRa1* expression through the cell cycle, which may have confounded earlier results that relied on non-quantitative RT-PCR assays (data not shown). We cannot exclude the possibility that differences between *SIR3-EBD* and earlier inducible alleles contributed to the different results. Indeed, Lazarus & Holmes’s use of the galactose promoter to drive *SIR3* expression would increase Sir3 dosage above wild-type levels, and alter the stoichiometry of the Sir protein complex. We expect Sir protein concentration and stoichiometry to be important parameters in regulating silencing establishment.

### Euchromatic histone mark removal was a key cell-cycle-regulated step in silencing establishment

Removal of Dot1-deposited methylation of H3K79 was a critical step in silencing establishment. This finding was consistent with earlier studies of cycling cells, which found that *dot1*Δ cells established silencing more quickly than wild-type cells (Katan-Khaykovich and Struhl, 2005; Osborne et al., 2009). Removal of this mark appears to be the primary reason why cells need to progress through S phase to establish silencing. Dot1 is thought to reduce Sir3 affinity for the nucleosome core by methylating H3K79 (Martino et al., 2009; Ng et al., 2002a). In addition to Dot1 and Sir3 both binding the nucleosome core at H3K79, they also compete for binding to the H4 tail, and deacetylation of the tail by Sir2 is thought to favor Sir3 binding at the expense of Dot1(Altaf et al., 2007). Thus, through modifications at H4K16 and H3K79, transcription and silencing mutually antagonize each other. In a cell in G1, even if Sir2/3/4 are able to displace Dot1 by deacetylating H4K16, H3K79me will remain until histones are turned over, which seems to explain the S-phase requirement for silencing establishment.

Silencing establishment at *HMR* does not require replication of the locus, as shown by the ability of excised episomes bearing *HMR* but no replication origin to establish silencing in an S-phase-dependent manner (Kirchmaier and Rine, 2001; Li et al., 2001). This finding presented a major mystery: what S-phase-specific process other than replication fork passage drives silencing establishment? Our results suggest that an influx of H3 molecules lacking methylation at K79 could be the solution. Replication-independent histone exchange can occur throughout the cell cycle (Dion et al., 2007; Rufiange et al., 2007; Schlissel and Rine, 2019), which means that a replicating or non-replicating copy of *HMR* can incorporate histone molecules from the nuclear pool. Outside of S phase, ∼90% of all H3 in the nucleus is methylated at K79 (Van Leeuwen et al., 2002), so histone exchange would likely lead to incorporation of the silencing-refractory methylated form. However, during S phase, a large quantity of newly-synthesized non-methylated H3 is present. Therefore, histone incorporation during S phase through either replication-coupled chromatin assembly or replication-independent histone would lead to incorporation of many H3 molecules that are not methylated at K79. This might explain why silencing establishment can occur at *HMR*, whether it is replicated or not, and why that establishment depends on S phase.

We found that *H3K79R* mutants, which mimicked the non-methylated state of H3K79, also established silencing in G1-arrested cells and did so even more strongly than *dot1*Δ mutants. A simple explanation for this difference in impact of the two mutations could be that Sir3 binds more strongly to arginine than lysine at position 79. Alternatively, Dot1 has been shown to have several methyltransferase-independent functions, and it was possible that one of these functions acted to promote silencing. In particular, Dot1 has recently been shown to possess histone chaperone activity that is independent of its ability to methylate histones (Lee et al., 2018). In addition, Dot1 has the methyltransferase-independent ability to stimulate ubiquitination of histone H2B (van Welsem et al., 2018). The latter result is particularly interesting, because H2B ubiquitination is itself required for both H3K79 methylation (Briggs et al., 2002; Ng et al., 2002b) and H3K4 methylation (Dover et al., 2002; Sun and Allis, 2002). Conflicting reports have pointed to a role of H3K4 methylation in silencing (Fingerman et al., 2005; Mueller et al., 2006; Santos-Rosa et al., 2004). Thus, it is possible that in a *dot1*Δ mutant, the removal of H3K79me *per se* promotes silencing, but an indirect effect through H2Bub and/or H3K4me partially counteracts the H3K79me effect.

The histone acetyltransferases Sas2 and Rtt109 also had roles in limiting silencing establishment to S phase. Individually, *sas2*Δ and *rtt109*Δ mutations led to partial silencing establishment in G1-arrested cells, and each of these effects was additive with a *dot1*Δ mutation. Acetylation of H4K16 by Sas2, like methylation of H3K79 by Dot1, is critical in distinguishing euchromatin and heterochromatin. Interestingly, in a previous study, while *dot1*Δ sped silencing establishment at *HML*, *sas2*Δ delayed silencing establishment by that assay (Osborne et al., 2009). The single-cell α-factor response assay used in that study required cells to fully repress *HML* to gain the **a** mating type identity, whereas our assay used more direct measures of changes in transcription at *HML* and *HMR*. Thus, one explanation consistent with both results is that *sas2*Δ cells begin silencing more readily than wild-type cells, but take more cell cycles to reach full repression. This could be the result of the competing effects of the *sas2*Δ mutation: hypoacetylation of histones at *HML* and *HMR* might increase Sir protein recruitment, while the global pool of hypoacetylated histones can also titrate Sir proteins away from *HML* and *HMR*.

The ability of *rtt109*Δ cells to drive partial silencing establishment in G1-arrested cells was surprising. Like Sas2, Rtt109 binds to Asf1 and acetylates newly-synthesized histones (Driscoll et al., 2007), but H3K56 acetylation is removed after S phase by the sirtuins Hst3 and Hst4 (Celic et al., 2006). The residual H3K56ac present outside of S phase is due to transcription-coupled histone turnover, which incorporates new histones marked with H3K56ac (Rufiange et al., 2007). A negative role for H3K56 acetylation in silencing has been observed, although this has not been well-characterized (Dodson and Rine, 2015; Miller et al., 2008). One simple model is that H3K56ac favors transcription, and thus impedes silencing establishment. However, given genome-wide acetylation and deacetylation of H3K56, indirect effects cannot be excluded.

Do the contributions of *DOT1*, *SAS2*, and *RTT109* completely resolve the cell-cycle requirement for silencing establishment? In *dot1*Δ mutants, S phase still dramatically accelerated silencing establishment, indicating that some feature of S phase beyond H3K79me removal was important in those cells. In addition, we found no case in which silencing establishment in G1-arrested cells matched the degree of silencing observed after overnight growth in estradiol. However, the ∼90% repression observed in, e.g., G1-arrested *H3K79R* cells should be sufficient to completely turn off transcription at *HML* and *HMR* in the majority of cells (see **Figure 2**). The quantitative gap in the level of silencing seen at steady state and that which is achieved in the experiments reported here could reflect a requirement for further cell-cycle steps or more time to complete silencing establishment. Others have identified a cell-cycle window between G2/M and G1 that contributes to silencing establishment (Lau et al., 2002), and none of our data were inconsistent with that result. Identifying mutant conditions in which G1-arrested cells and cycling cells establish silencing at an equal rate will be required before the cell-cycle regulated establishment of silencing is fully understood.

Together, our data suggest that silencing establishment cannot proceed without removal of histone modifications that favor transcription. In this view, at any stage of the cell cycle, Sir proteins can bind to *HML* and *HMR*. Passage through S phase leads to incorporation of new histones, which, crucially, lack H3K79 methylation. This decrease of H3K79me by half leads to both further Sir binding and decreased transcription. However, one cell cycle is not sufficient to fully deplete activating marks, and successive passages through S phase complete the process of silencing establishment.

## MATERIALS AND METHODS

### Yeast strains

Strains used in this study are listed in **Table S1**. All strains were derived from the W303 background using standard genetic techniques (Dunham et al., 2015; Gietz and Schiestl, 2007). Deletions were generated using one-step replacement with marker cassettes (Goldstein and McCusker, 1999; Gueldener, 2002). The tRNA gene *tT(AGU)C* was seamlessly deleted using the “delitto perfetto” technique as described previously (Storici and Resnick, 2006). The *MCD1-AID* strain was generated by first inserting *O.s.TIR1* at the *HIS3* locus by transforming cells with PmeI-digested pTIR2 (Eng et al., 2014). Then, *3xV5-AID2:KanMX* was amplified from pVG497 (a gift from Vincent Guacci and Douglas Koshland) with primers that included homology to *MCD1*, followed by transformation. The mutant allele *HML\** was synthesized as a DNA gene block (Integrated DNA Technologies) and integrated using CRISPR-Cas9 technology as previously described (Brothers and Rine, 2019). The *EBD* sequence was amplified by PCR from *cre-EBD78* in the strain UCC5181 (Lindstrom and Gottschling, 2009) with primers that included homology to *SIR3,* then transformed using CRISPR-Cas9. Mutations of *HHT1* and *HHT2* were generated using CRISPR-Cas9, with oligonucleotide repair templates. For all mutant analyses, at least two independent transformants or meiotic segregants were tested.

### Culture growth and cell-cycle manipulations

All experiments were performed on cells growing in yeast extract peptone + 2% dextrose (YPD) at 37°, which led to more switch-like behavior for *SIR3-EBD* than growth at 30°. For steady-state measurements, cells were grown overnight in YPD, then diluted and grown in fresh YPD to a density of ∼2-8 × 10^6^ cells/mL. For cell-cycle control experiments, cells were grown overnight in YPD, followed by dilution and growth in fresh YPD for ≥ 3 hours until cultures reached a density of ∼2 × 10^6^ cells/mL. Then, α factor (synthesized by Elim Biopharmaceuticals; Hayward, CA) was added to a final concentration of 10 nM and the cultures were incubated for ∼2 hours until >90% of cells appeared unbudded. For *rtt109*Δ strains, this incubation was ∼3 hours. For experiments with prolonged α-factor arrests, additional α factor was added every ∼2 hours to maintain the arrest. To release cells from α-factor arrest, protease from *Streptomyces griseus* (Sigma-Aldrich P5147; St. Louis, MO) was added to the media at a final concentrat1io0n ofn 0.1 mg/mL. To re-arrest cells at G2/M, nocodazole (Sigma-Aldrich M1404) was added to a final concentration of 15 μg/mL. For *SIR3-EBD* induction, β-estradiol (Sigma-Aldrich E8875) was added to a final concentration of 50 μM from a 10 mM stock in ethanol. For Mcd1-AID depletion, 3-indoleacetic acid (auxin; Sigma-Aldrich I2886) was added to a final concentration of 750 μM from a 1 M DMSO stock.

### RNA extraction and RT-qPCR

For each sample, at least ∼1 × 10^7^ cells were collected by centrifugation and RNA was purified using an RNeasy Mini Kit (Qiagen 74104; Hilden, Germany), including on-column DNase digestion (Cat No. 79254), according to manufacturer’s recommendations. 2 μg of RNA was reverse transcribed using SuperScript III reverse transcriptase (Thermo Fisher Scientific 18080044; Waltham, MA) and an “anchored” oligo-dT primer (an equimolar mixture of primers with the sequence T_20_VN, where V represents any non-T base). qPCR was performed using the DyNAmo HS SYBR Green qPCR kit (Thermo Fisher Scientific F410L), including a Uracil-DNA Glycosylase (Thermo Fisher Scientific EN0362) treatment, and samples were run using an Agilent Mx3000P thermocycler. Oligonucleotides used for qPCR are listed in **Table S2**. cDNA abundance was calculated using a standard curve and normalized to the reference gene *ALG9*. Each reaction was performed in triplicate, and a matched non-reverse-transcribed sample was included for each sample.

### Chromatin immunoprecipitation and sequencing

For ChIP experiments, ∼5 × 10^8^ cells were crosslinked in 1% formaldehyde at room temperature for 60 minutes (**Figure 3A, 3B**) or 15 minutes (all other figures). Following a 5-minute quench in 300 mM glycine, cells were washed twice in ice-cold TBS and twice in ice-cold FA lysis buffer (50 mM HEPES, pH 7.5; 150 mM NaCl, 1 mM EDTA, 1% Triton, 0.1% sodium deoxycholate) + 0.1% SDS + protease inhibitors (cOmplete EDTA-free protease inhibitor cocktail, Sigma-Aldrich 11873580001). Cell pellets were then either flash frozen or lysed. For lysis, cell pellets were resuspended in 1 mL FA lysis buffer without EDTA + 0.1% SDS and ∼500 μL 0.5-mm zirconia/Silica beads (BioSpec Products; Bartlesville, OK) were added. Cells were lysed using a FastPrep-24 5G (MP Biomedicals; Irvine, CA) with 6.0 m/s beating for 20 seconds followed by 2 minutes on ice, repeated 4 times total. Lysate was transferred to a new microcentrifuge tube, then spun at 4° for 15 minutes at 17,000 rcf. The pellet was resuspended in 1 mL FA lysis buffer without EDTA + 0.1% SDS + 2 mM CaCl_2_. The samples were incubated at 37° for 5 minutes, followed by addition of 4 U MNase (Worthington Biochemical LS004798; Lakewood, NJ) per 1 × 10^7^ cells. Samples were placed on an end-over-end rotator at 37° for 20 minutes, followed by addition of 1.25 mM EDTA to quench the digestion. Samples were spun at 4° for 10 minutes at 17,000 rcf, and the supernatant containing fragmented chromatin was saved.

The fragmented chromatin was split, saving 50 μL as the input sample, and using the remaining ∼900 μL for immunoprecipitation. Immunoprecipitation for Sir4-13xMyc was performed using 50 μL Pierce Anti-c-myc magnetic beads (Thermo Fisher Scientific 88843) per sample. Beads were equilibrated by washing 5x in FA Lysis buffer + 0.1% SDS + 0.05% Tween, then resuspended in in 50 μL per sample of FA Lysis buffer + 0.1% SDS + 0.05% Tween. 50 μL beads and 0.5 mg/mL BSA (NEB B9000S; Ipswich, MA) were added to each fragmented chromatin sample, and placed on an end-over-end rotator at 4° overnight. The following washes were performed, placing each sample on an end-over-end rotator for ∼5 minutes between each: 2x washes with FA Lysis + 0.1% SDS + 0.05% Tween; 2x washes with Wash Buffer #1 (FA Lysis buffer + 0.25 M NaCl + 0.1% SDS + 0.05% Tween); 2x washes with Wash Buffer #2 (10 mM Tris, pH 8; 0.25 M LiCl; 0.5% NP-40; 0.5% sodium deoxycholate; 1 mM EDTA + 0.1% SDS + 0.05% Tween); and 1x wash with TE + 0.05% Tween. Samples were eluted by adding 100 μL TE + 1% SDS to the beads and incubating at 65° for 10 minutes with gentle shaking. The eluate was saved, and the elution was repeated for a total eluate volume of 200 μL. Input samples were brought to a total volume of 200 μL with TE + 1% SDS. IP and input samples were incubated with 10 μL of 800 U/mL Proteinase K (NEB P8107S) at 37° for 2 hours, followed by overnight incubation at 65° to reverse crosslinking.

DNA was purified using a QIAquick PCR purification kit (Qiagen 28104). Libraries were prepared for high-throughput sequencing using the Ultra II DNA Library Prep kit (NEB E7645L). For IP samples, the entire purified sample was used in the library prep reaction. For the input samples, 10 ng were used. Samples were multiplexed and paired-end sequencing was performed using either a MiniSeq or NovaSeq 6000 (Illumina; San Diego, CA).

Sequencing reads were aligned using Bowite2 (Langmead and Salzberg, 2012) to a reference genome derived from SacCer3, modified to include the mutant *HML\** and *mat*Δ. Aligned reads corresponding to monoucleosomes (130-180 bp) were normalized to the non-heterochromatic genome-wide median (i.e., to the genome-wide median excluding rDNA, subtelomeric regions, and all of chromosome III). Analysis was performed using custom Python scripts and displayed using IGV (Thorvaldsdóttir et al., 2013). Reads were deposited at xxxxx.

### Single-molecule RNA fluorescence in situ hybridization

For smRNA-FISH experiments, cells were grown and cell-cycle arrests were performed as described above. Preparation of samples for imaging was as previously described (Chen et al., 2018). The only modification from that protocol was the concentration of zymolyase used: for cycling cells, 5 μL of 1.25 mg/mL zymolyase-100T (VWR IC320932; Radnor, PA) was used during the spheroplasting; for arrested cells, 5 μL of 2.5 mg/mL zymolyase-100T was used. Probes were synthesized by Stellaris (Biosearch Technologies; Novato, CA). Probes for *cre* and *KAP104* were previously described (Dodson and Rine, 2015). The sequences of all probes, including newly-designed probes for *a1* are listed in **Table S3**. Probes for *cre* and *a1* were coupled to Quasar 670 dye, while probes for *KAP104* were coupled to CAL Fluor Red 590 dye. Probes for *cre* and *KAP104* were used at a final concentration of 100 nM. Probes for *a1* were used at a final concentration of 25 nM.

Imaging was performed on an Axio Observer Z1 inverted microscope (Zeiss; Oberkochen, Germany) equipped with a Plan-Apochromat 63x oil-immersion objective (Zeiss), pE-300 ultra illumination system (CoolLED; Andover, UK), MS-2000 XYZ automated stage (Applied Scientific Instrumentation; Eugene, OR), and 95B sCMOS camera (Teledyne Photometrics; Tucson, AZ). The following filter sets were used: for Quasar 670, Cy5 Narrow Excitation (Chroma Cat No. 49009; Bellows Falls, VT); for CAL Fluor Red 590, filter set 43 HE (Zeiss); for DAPI, multiband 405/488/594 filter set (Semrock Part No. LF405/488/594-A-000; Rochester, NY). The microscope was controlled using Micro-manager software (Edelstein et al., 2014). Z-stack images were taken with a total height of 10 μm and a step size of 0.2 μm. Manual cell outlining and automatic spot detection was performed using FISH-quant (Mueller et al., 2013) and data was plotted using ggplot2 (Wickham, 2016). Representative images were generated using FIJI (Schindelin et al., 2012).

### Protein extraction and immunoblotting

Protein extraction and immunoblotting were performed as previously described (Brothers and Rine, 2019). The primary antibodies used were 1:20,000 Rabbit anti-Hexokinase (Rockland #100-4159; Limerick, PA) and 1:2,500 Mouse anti-V5 (Thermo Fisher Scientific R960-25). The secondary antibodies used were 1:20,000 IRDye 800CW Goat anti-Mouse (Li-Cor; Lincoln, NE) and 1:20,000 IRDye 680RD Goat anti-Rabbit (Li-Cor).

### Flow cytometry

For every cell-cycle experiment, an aliquot of cells was taken for flow-cytometry analysis of DNA content to monitor cell-cycle stage. Sample preparation and flow cytometry was performed as described previously (Schlissel and Rine, 2019). A representative sample of flow cytometry profiles is shown in **Figure 5—figure supplement 1**.

## Supporting information

Combined_tables

## ACKNOWLEDGMENTS

We are grateful to all members of the Rine laboratory and Koshland laboratory for helpful discussions and comments on this work. We thank Vincent Guacci for providing plasmids and for his valuable guidance, along with Lorenzo Costantino, on performing cell-cycle manipulations. We thank Itamar Patek for his help with strain construction. We give special thanks to Marc Fouet for his generous assistance with microscopy. This work relied on the Vincent J. Coates Genomics Sequencing Laboratory at UC Berkeley, supported by NIH S10 OD018174 Instrumentation Grant. This work was funded by grants from the National Institutes of Health to J.R. (GM31105, GM120374). D.G. received support from NIH training grants (T32 GM007127, T32 HG000047) and a National Science Foundation Graduate Research Fellowship (Grant No. 1752814).

---

**Figure 1—figure supplement 1:**
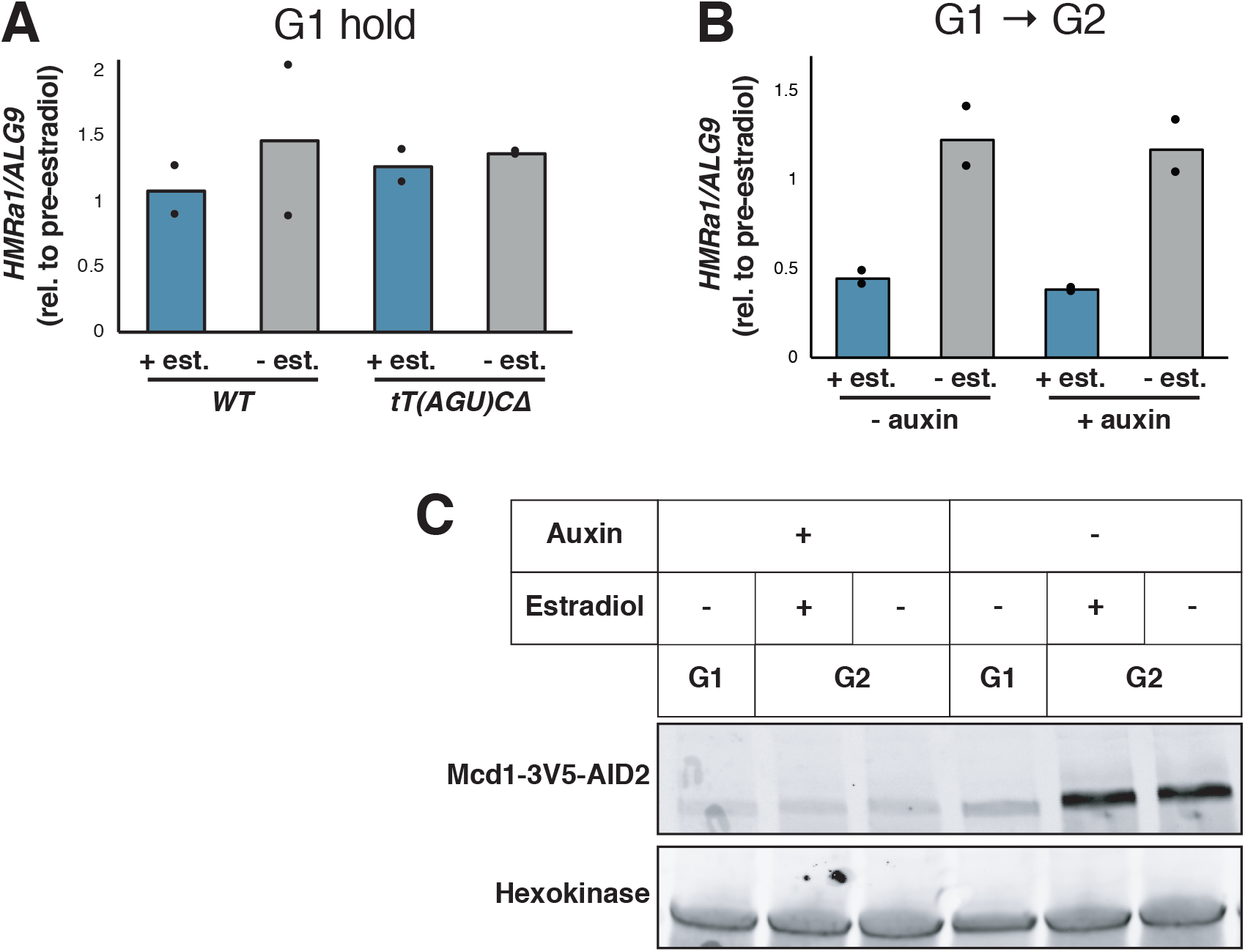
Effects of cohesin depletion and *tT(AGU)C* deletion on silencing establishment. **(A)** *SIR3-EBD* strains (JRY12269, JRY12270) and *SIR3-EBD* strains with seamless deletion of *tT(AGU)C* (JRY12267; JRY12268) were arrested in G1 with α factor, then split, with half the culture receiving estradiol and half receiving ethanol. Samples were collected after 3 hours for RT-qPCR, with each sample normalized to its own pre-estradiol value. **(B)** Cells with *MCD1-AID* (JRY12560, JRY12561) were arrested in G1 with α factor, then split, with half receiving auxin and the other half receiving DMSO (solvent control). After 30 minutes, each culture was further split, with half receiving estradiol and the other half ethanol. All cultures were released to G2/M by addition of protease and nocodazole. Cells were collected after 3 hours for RT-qPCR, with each sample normalized to its own pre-estradiol value. **(C)** Immunoblot analysis showing Mcd1-AID depletion for experiment described in (B).

**Figure 1—figure supplement 2:**
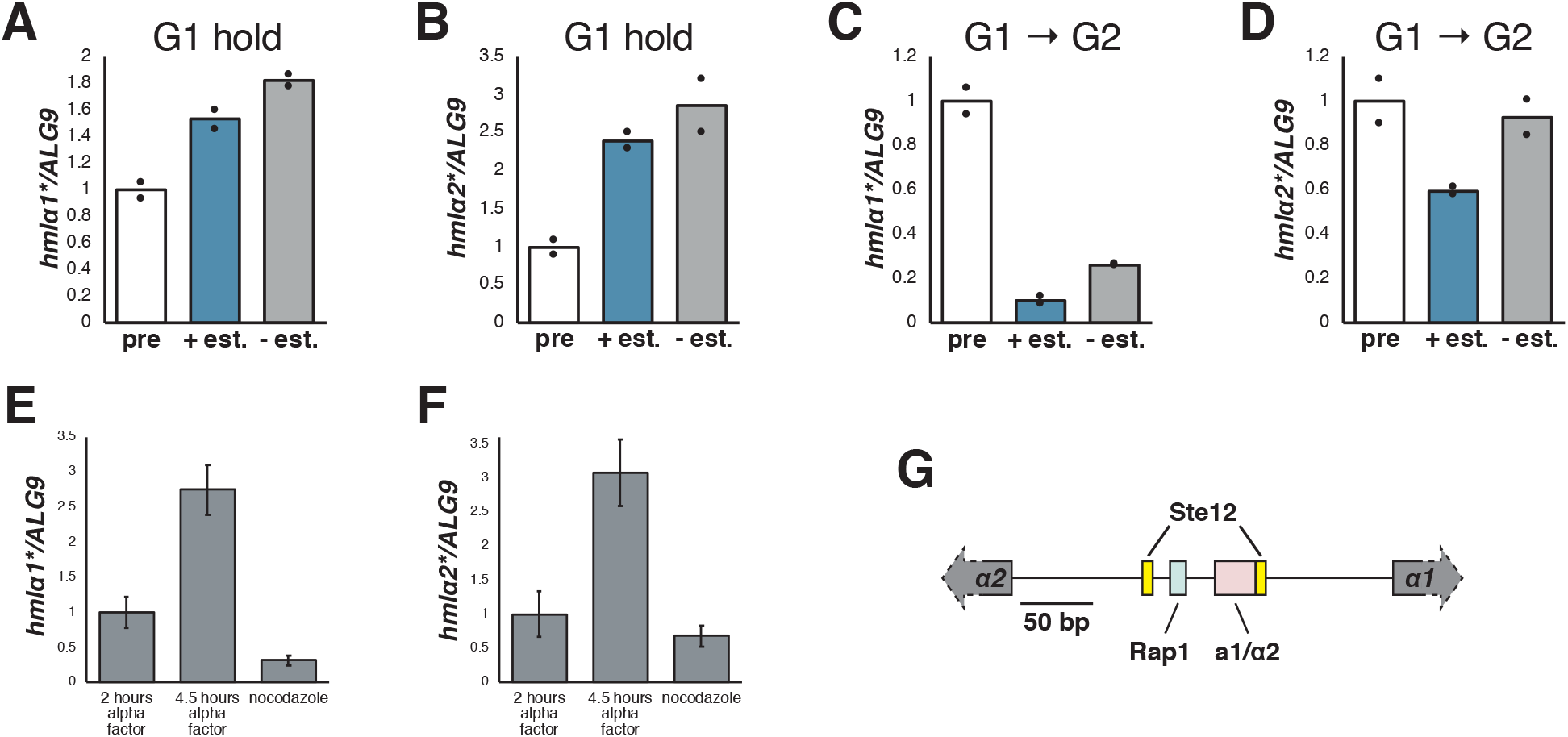
Silencing establishment required S phase at *HML\**. **(A)** Cells with *SIR3-EBD* (JRY12169) were arrested in G1 with α factor, then split, with one sub-culture receiving estradiol and the other receiving ethanol. Cells were collected after 6 hours in estradiol and analyzed by RT-qPCR for *hmlα1\**. **(B)** RT-qPCR for *hmlα2\** in the same cells described in (A). **(C)** Cells were arrested in G1 with α factor, then split and released to G2/M with protease and nocodazole, with one sub-culture receiving estradiol and the other receiving ethanol. Cells were collected after 3 hours in estradiol and analyzed by RT-qPCR for *hmlα1\**. The pre-estradiol samples for this experiment were the same as those described in (A). **(D)** RT-qPCR for *hmlα2\** in the same cells described in (C). **(E)** Cells lacking *SIR3* (JRY11966) were arrested in α factor for 2 hours, then split, with half staying in α factor and the other being released to G2/M by addition of protease and nocodazole. Cells were analyzed by RT-qPCR for *hmlα1*,* with error bars representing standard deviation among technical replicates (N=1 biological sample). **(F)** RT-qPCR for *hmlα2\** in the cells described in (E). **(G)** Map of the *α1/α2* promoter, showing newly-identified Ste12 motifs (TGAAACA) along with previously-identified binding sites for Rap1 and a1/α2.

**Figure 2—figure supplement 1:**
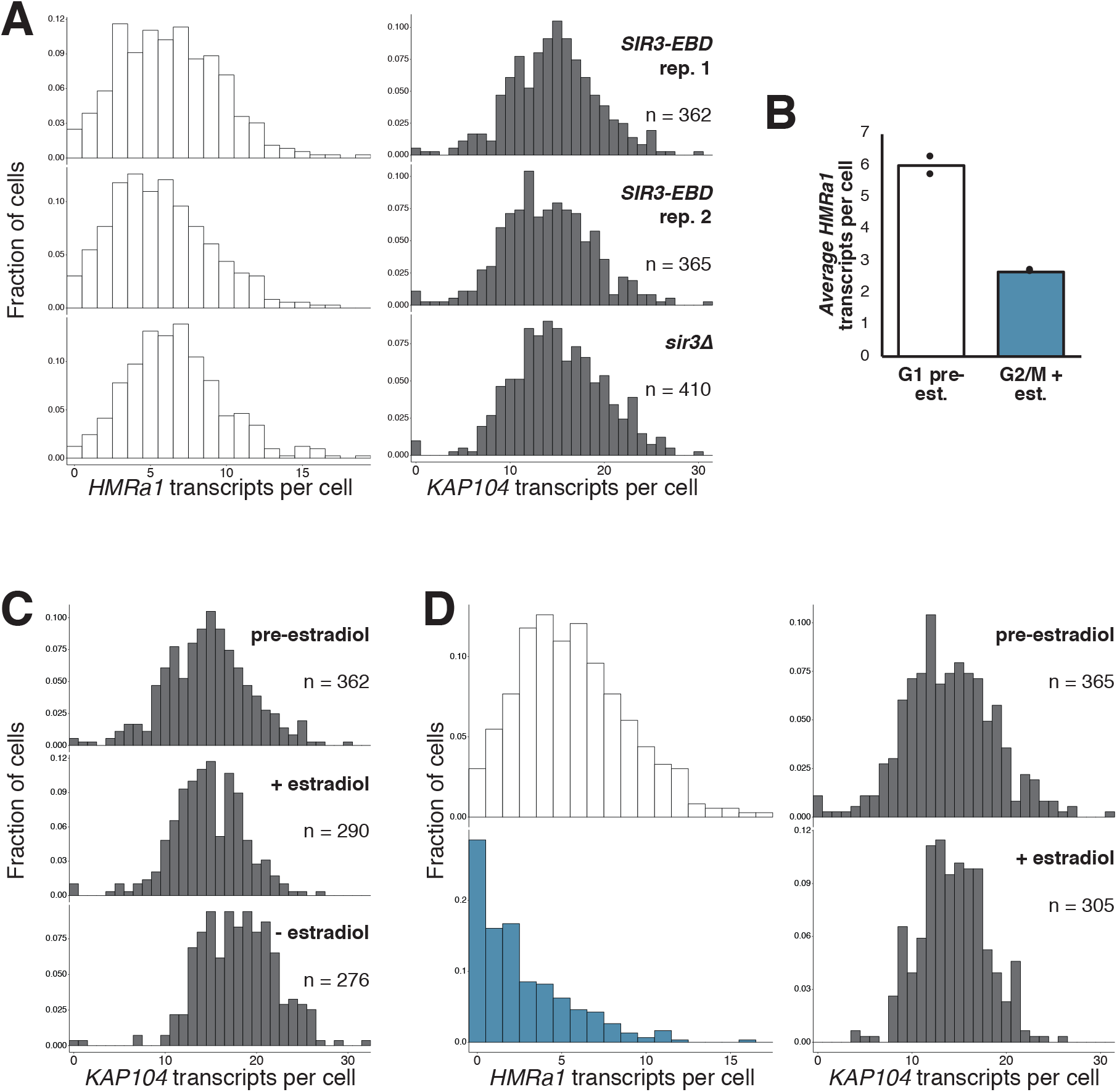
Gradual silencing establishment at *HMR*. **(A)** Quantification of *HMRa1* and *KAP104* transcripts in G1-arrested *SIR3-EBD* (JRY11762, JRY11763) and *sir3*Δ (JRY11966) cells. The *SIR3-EBD* data are the same as displayed in Figure 2B and Figure 2—figure supplement 1D. **(B)** Average number of *HMRa1* transcripts per cell before and after silencing establishment, quantified from data shown in Figure 2B and Figure 2—figure supplement 1D. Compare changes in mRNA levels to values from bulk measurement in Figure 1E. **(C)** Quantification of *KAP104* transcripts per cell for experiment described in Figure 2B. **(D)** Replicate experiment to that shown in Figure 2B with isogenic cells (JRY11763).

**Figure 2—figure supplement 2:**
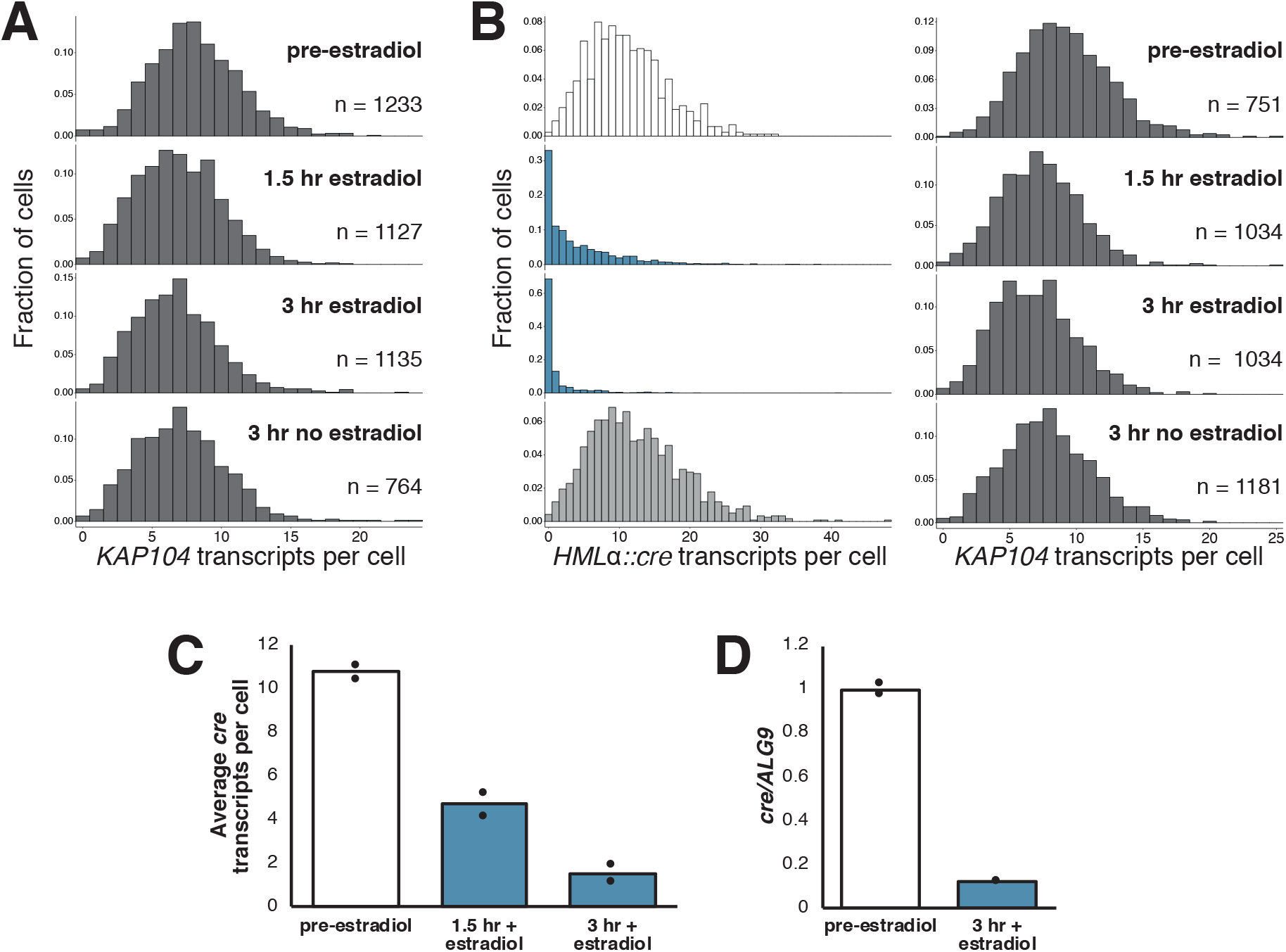
Gradual silencing establishment at *HMLα∷cre*. **(A)** Quantification of *KAP104* transcripts per cell for experiment described in Figure 2C. **(B)** Replicate experiment to that shown in **Figure 2B** with isogenic cells (JRY12513). **(C)** Average number of *cre* transcripts per cell before and during silencing establishment, quantified from data shown in Figure 2C and Figure 2—figure supplement 2B. **(D)** Bulk measurement RT-qPCR for *cre* during silencing establishment in an analogous experiment to that described in Figure 2C.

**Figure 3—figure supplement 1:**
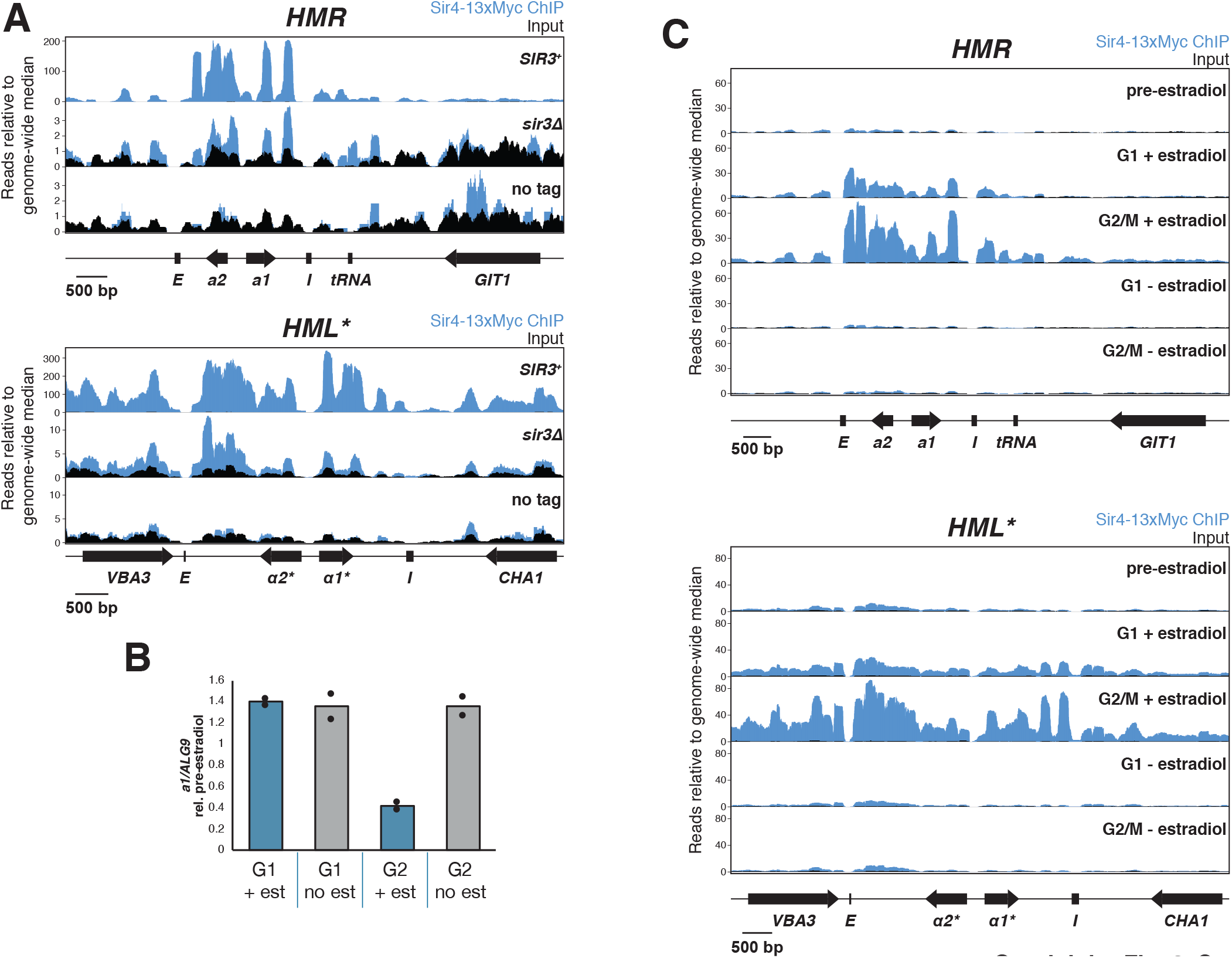
Silencing establishment ChIP-seq. All ChIP-seq panels show Sir4-13xMyc in blue and input in black. Read counts were normalized to the non-heterochromatin genome-wide median. IP and input values are plotted on the same scale. **(A)** ChIP-seq for Sir4-13xMyc at *HMR* (top) and *HML\** (bottom) in strains with *SIR3^+^* (JRY12171), *sir3*Δ (JRY12167). Also shown is an equivalent experiment in cells with untagged Sir4 (JRY12269). Cells were grown to mid-log phase and fixed for 15 minutes in formaldehyde. **(B)** RT-qPCR analysis of silencing establishment at *HMRa1* from samples that were used for ChIP-seq described in Figure 3C and Figure 3—figure supplement 1C. Values are scaled to the pre-estradiol value for each sample. **(C)** Replicate experiment to that described in Figure 3C, using an isogenic strain (JRY12170).

**Figure 4—figure supplement 1:**
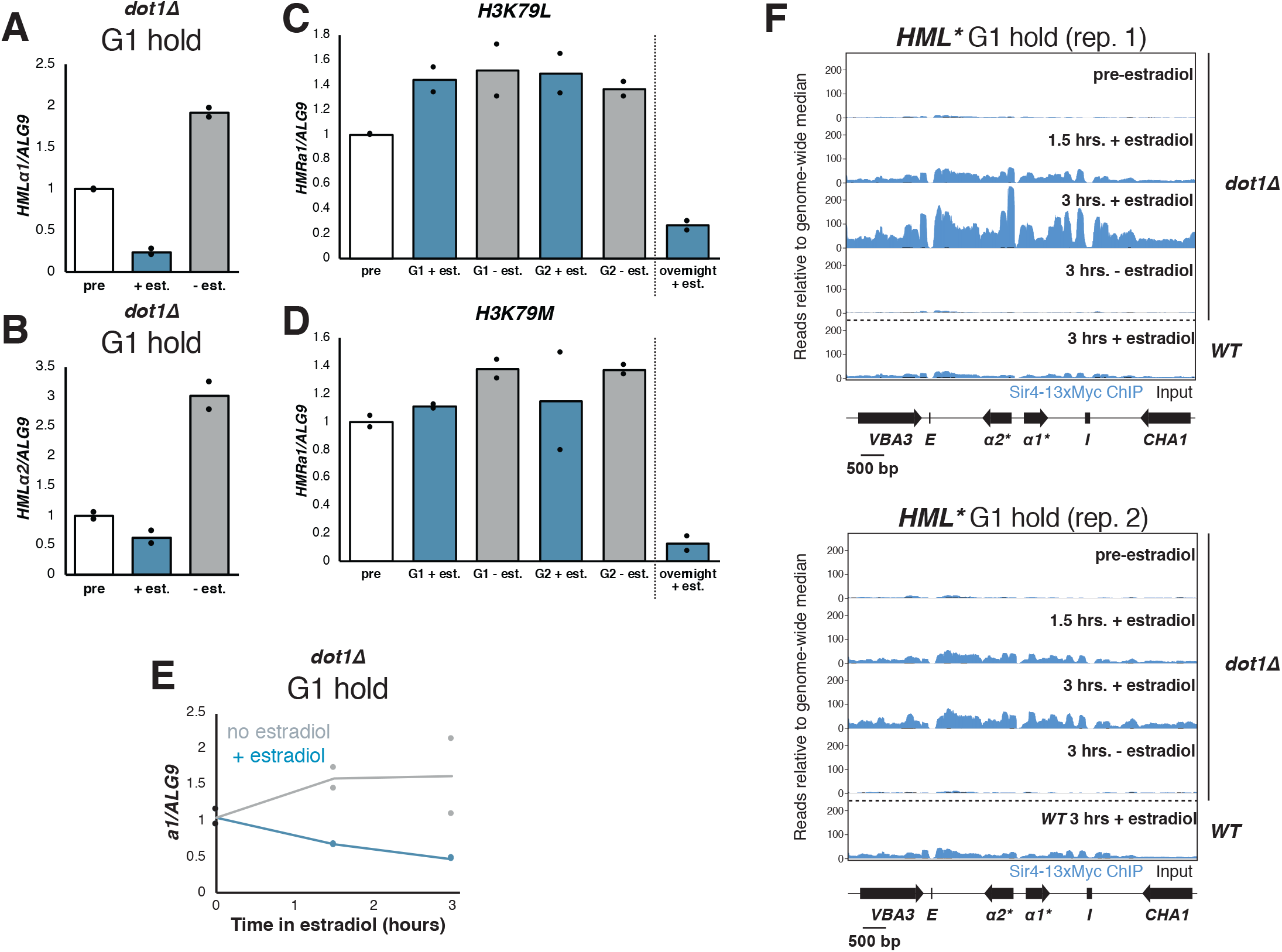
Silencing establishment in *dot1*Δ cells. **(A)** *dot1*Δ cells (JRY12443) were arrested in G1 with α factor, then split, with one sub-culture receiving estradiol and the other receiving ethanol. Cells were collected after 6 hours in estradiol and RT-qPCR was performed for *hmlα1\**. The data in this plot are also shown in Figure 5B for comparison with other mutants. **(B)** RT-qPCR for *hmlα2\** in the cells described in (A). **(C)** Cells with lysine 79 of H3 mutated to leucine in both *HHT1* and *HHT2* in two isogenic strains (*H3K79L;* JRY12854, JRY12855) were arrested in G1 with α factor, then split four ways. Two cultures were kept in G1, with one receiving estradiol and the other ethanol. The other two cultures were released to G2/M by addition of protease and nocodazole, and either estradiol or ethanol. Cells were collected after 3 hours for the G2/M samples and after 6 hours for the G1 samples, and RT-qPCR was performed for *HMRa1*. Also shown is a sample grown overnight in medium with estradiol. **(D)** Cells with lysine 79 of H3 mutated to methionine in both *HHT1* and *HHT2* in two isogenic strains (*H3K79M*; JRY12857, JRY12858) were subjected to the same experiment described in (C). **(E)** RT-qPCR analysis of silencing establishment at *HMRa1* from samples that were used for ChIP-seq shown in Figure 4E and Figure 4—figure supplement 1F. **(F)** Sir4-13xMyc binding at *HML\** from the same samples displayed in Figure 4E. Sir4-13xMyc is in blue and input in black, plotted on the same scale. Read counts were normalized to the non-heterochromatin genome-wide median.

**Figure 5—figure supplement 1:**
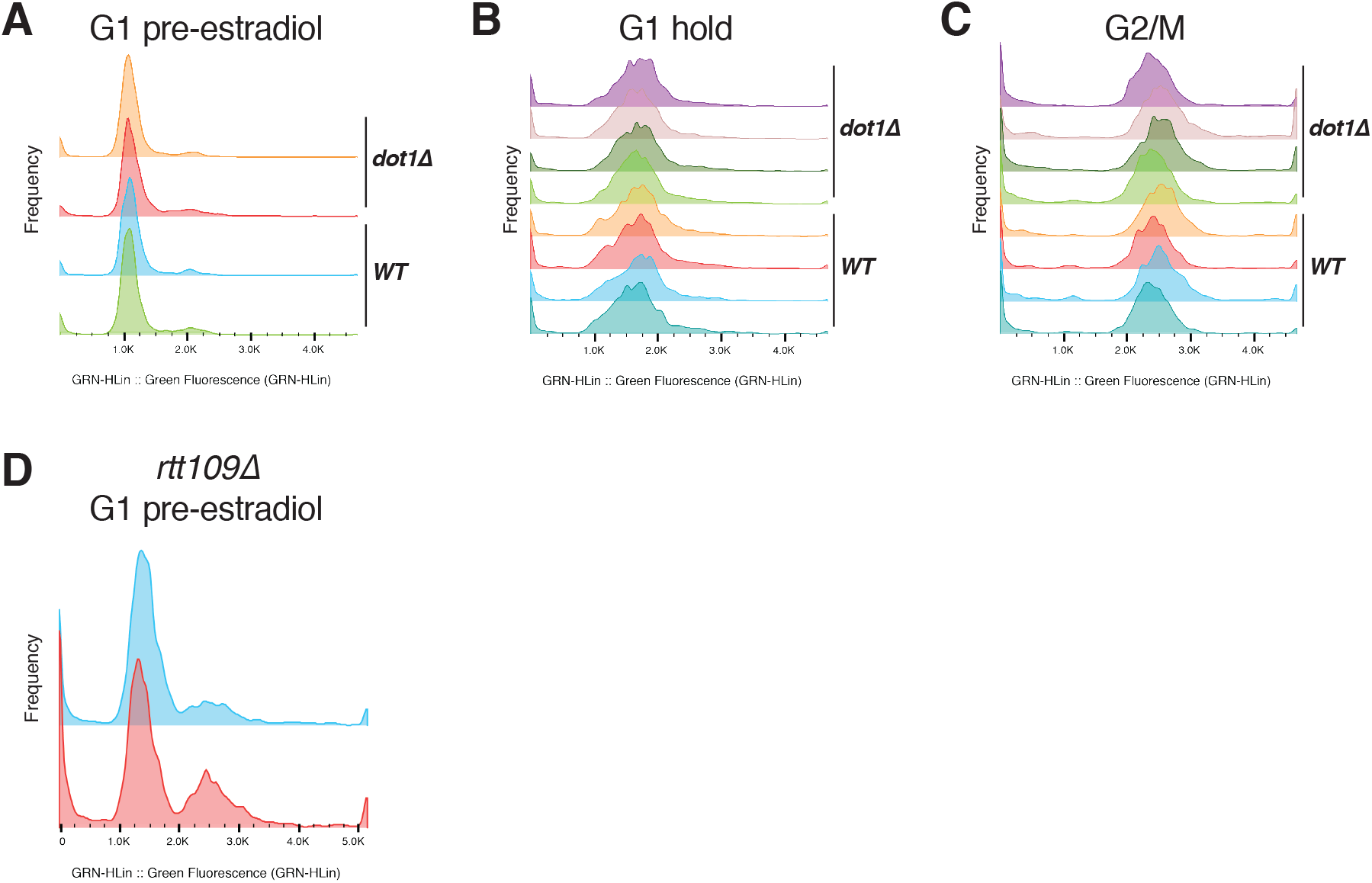
Representative flow cytometry profiles. **(A)** Wild-type (JRY12169) and *dot1*Δ (JRY12443) cells arrested in G1 with α factor for ∼2 hours. **(B)** Wild-type and *dot1*Δ cells kept in G1 for 6 hours, as in, e.g., Figure 1D. **(C)** Wild-type and *dot1*Δ cells after 3 hours in G2/M, as in, e.g., Figure 1E. **(D)** Cells lacking *RTT109* (JRY12689, JRY12691) arrested in G1 with α factor for ∼3 hours did not arrest uniformly.

**Table S1: Yeast strains used in this study.** All strains listed were generated for this study and derived from the W303 background.

**Table S2: Oligonucleotides used for RT-qPCR**

**Table S3: Probes used for smRNA-FISH**

## REFERENCES

Adkins MW, Carson JJ, English CM, Ramey CJ, Tyler JK. 2007. The histone chaperone anti-silencing function 1 stimulates the acetylation of newly synthesized histone H3 in S-phase. J Biol Chem 282:1334–1340. doi:10.1074/jbc.M608025200

Altaf M, Utley RT, Lacoste N, Tan S, Briggs SD, Côté J. 2007. Interplay of Chromatin Modifiers on a Short Basic Patch of Histone H4 Tail Defines the Boundary of Telomeric Heterochromatin. Mol Cell 28:1002–1014. doi:10.1016/j.molcel.2007.12.002

Armache K-J, Garlick JD, Canzio D, Narlikar GJ, Kingston RE. 2011. Structural basis of silencing: Sir3 BAH domain in complex with a nucleosome at 3.0 Å resolution. Science 334:977–82. doi:10.1126/science.1210915

Briggs SD, Xiao T, Sun Z-W, Caldwell JA, Shabanowitz J, Hunt DF, Allis CD, Strahl BD. 2002. Trans-histone regulatory pathway in chromatin. Nature 418:498–498. doi:10.1038/nature00970

Brothers M, Rine J. 2019. Mutations in the PCNA DNA Polymerase Clamp of Saccharomyces cerevisiae Reveal Complexities of the Cell Cycle and Ploidy on Heterochromatin Assembly. Genetics 213:449–463. doi:10.1534/genetics.119.302452

Celic I, Masumoto H, Griffith WP, Meluh P, Cotter RJ, Boeke JD, Verreault A. 2006. The Sirtuins Hst3 and Hst4p Preserve Genome Integrity by Controlling Histone H3 Lysine 56 Deacetylation. Curr Biol 16:1280–1289. doi:10.1016/j.cub.2006.06.023

Chen J, McSwiggen D, Ünal E. 2018. Single molecule fluorescence in situ hybridization (SmFISH) analysis in budding yeast vegetative growth and meiosis. J Vis Exp 2018:1–14. doi:10.3791/57774

De Vos D, Frederiks F, Terweij M, Van Welsem T, Verzijlbergen KF, Iachina E, De Graaf EL, Maarten Altelaar AF, Oudgenoeg G, Heck AJR, Krijgsveld J, Bakker BM, Van Leeuwen F. 2011. Progressive methylation of ageing histones by Dot1 functions as a timer. EMBO Rep 12:956–962. doi:10.1038/embor.2011.131

Dion MF, Kaplan T, Kim M, Buratowski S, Friedman N, Rando OJ. 2007. Dynamics of replication-independent histone turnover in budding yeast. Science 315:1405–8. doi:10.1126/science.1134053

Dodson AE, Rine J. 2015. Heritable capture of heterochromatin dynamics in Saccharomyces cerevisiae. Elife 4:e05007. doi:10.7554/eLife.05007

Dolan JW, Kirkman C, Fields S. 1989. The yeast STE12 protein binds to the DNA sequence mediating pheromone induction. Proc Natl Acad Sci 86:5703–5707. doi:10.1073/pnas.86.15.5703

Dover J, Schneider J, Tawiah-Boateng MA, Wood A, Dean K, Johnston M, Shilatifard A. 2002. Methylation of histone H3 by COMPASS requires ubiquitination of histone H2B by Rad6. J Biol Chem 277:28368–28371. doi:10.1074/jbc.C200348200

Driscoll R, Hudson A, Jackson SP. 2007. Yeast Rtt109 Promotes Genome Stability by Acetylating Histone H3 on Lysine 56. Science 315:649–652. doi:10.1126/science.1135862

Dunham M, Gartenberg M, Brown GW, editors. 2015. Methods in Yeast Genetics and Genomics: A Cold Spring Harbor Laboratory Course Manual, 2015 Edition. Cold Spring Harbor, NY: Cold Spring Harbor Laboratory Press.

Edelstein AD, Tsuchida MA, Amodaj N, Pinkard H, Vale RD, Stuurman N. 2014. Advanced methods of microscope control using μManager software. J Biol Methods 1:10. doi:10.14440/jbm.2014.36

Eng T, Guacci V, Koshland D. 2014. ROCC, a conserved region in cohesin’s Mcd1 subunit, is essential for the proper regulation of the maintenance of cohesion and establishment of condensation. Mol Biol Cell 25:2351–2364. doi:10.1091/mbc.E14-04-0929

Fillingham J, Recht J, Silva AC, Suter B, Emili A, Stagljar I, Krogan NJ, Allis CD, Keogh M-C, Greenblatt JF. 2008. Chaperone Control of the Activity and Specificity of the Histone H3 Acetyltransferase Rtt109. Mol Cell Biol 28:4342–4353. doi:10.1128/mcb.00182-08

Fingerman IM, Wu CL, Wilson BD, Briggs SD. 2005. Global loss of Set1-mediated H3 Lys4 trimethylation is associated with silencing defects in Saccharomyces cerevisiae. J Biol Chem 280:28761–28765. doi:10.1074/jbc.C500097200

Gartenberg MR, Smith JS. 2016. The Nuts and Bolts of Transcriptionally Silent Chromatin in Saccharomyces cerevisiae. Genetics 203:1563–1599. doi:10.1534/genetics.112.145243

Gietz RD, Schiestl RH. 2007. High-efficiency yeast transformation using the LiAc/SS carrier DNA/PEG method. Nat Protoc 2:31–34. doi:10.1038/nprot.2007.13

Goldstein AL, McCusker JH. 1999. Three new dominant drug resistance cassettes for gene disruption in Saccharomyces cerevisiae. Yeast 15:1541–1553. doi:10.1002/(SICI)1097-0061(199910)15:14<1541::AID-YEA476>3.0.CO;2-K

Gueldener U. 2002. A second set of loxP marker cassettes for Cre-mediated multiple gene knockouts in budding yeast. Nucleic Acids Res 30:23e–23. doi:10.1093/nar/30.6.e23

Hecht A, Laroche T, Strahl-Bolsinger S, Gasser SM, Grunstein M. 1995. Histone H3 and H4 N-termini interact with SIR3 and SIR4 proteins: a molecular model for the formation of heterochromatin in yeast. Cell 80:583–92. doi:10.1016/0092-8674(95)90512-x

Herskowitz I. 1989. A regulatory hierarchy for cell specialization in yeast. Nature 342:749–757. doi:10.1038/342749a0

Hoppe GJ, Tanny JC, Rudner AD, Gerber SA, Danaie S, Gygi SP, Moazed D. 2002. Steps in Assembly of Silent Chromatin in Yeast: Sir3-Independent Binding of a Sir2/Sir4 Complex to Silencers and Role for Sir2-Dependent Deacetylation. Mol Cell Biol 22:4167–4180. doi:10.1128/mcb.22.12.4167-4180.2002

Imai S, Armstrong CM, Kaeberlein M, Guarente L. 2000a. Transcriptional silencing and longevity protein Sir2 is an NAD-dependent histone deacetylase. Nature 403:795–800.

Imai S, Armstrong CM, Kaeberlein M, Guarente L. 2000b. Transcriptional silencing and longevity protein Sir2 is an NAD-dependent histone deacetylase. Nature 403:795–800. doi:10.1038/35001622

Johnson LM, Kayne PS, Kahn ES, Grunstein M. 1990. Genetic evidence for an interaction between SIR3 and histone H4 in the repression of the silent mating loci in Saccharomyces cerevisiae. Proc Natl Acad Sci U S A 87:6286–6290. doi:10.1073/pnas.87.16.6286

Katan-Khaykovich Y, Struhl K. 2005. Heterochromatin formation involves changes in histone modifications over multiple cell generations. EMBO J 24:2138–49. doi:10.1038/sj.emboj.7600692

Kimura A, Umehara T, Horikoshi M. 2002. Chromosomal gradient of histone acetylation established by Sas2p and Sir2p functions as a shield against gene silencing. Nat Genet 32:370–377. doi:10.1038/ng993

Kirchmaier AL, Rine J. 2006. Cell Cycle Requirements in Assembling Silent Chromatin in Saccharomyces cerevisiae. Mol Cell Biol 26:852–862. doi:10.1128/MCB.26.3.852-862.2006

Kirchmaier AL, Rine J. 2001. DNA replication-independent silencing in S. cerevisiae. Science 291:646–50. doi:10.1126/science.291.5504.646

Kitada T, Kuryan BG, Nga N, Tran H, Song C, Xue Y, Carey M, Grunstein M. 2012. Mechanism for epigenetic variegation of gene expression at yeast telomeric heterochromatin 2:2443–2455. doi:10.1101/gad.201095.112.is

Landry J, Sutton A, Tafrov ST, Heller RC, Stebbins J, Pillus L, Sternglanz R. 2000. The silencing protein SIR2 and its homologs are NAD-dependent protein deacetylases. Proc Natl Acad Sci 97:5807–5811. doi:10.1073/pnas.110148297

Langmead B, Salzberg SL. 2012. Fast gapped-read alignment with Bowtie 2. Nat Methods 9:357–359. doi:10.1038/nmeth.1923

Lau A, Blitzblau H, Bell SP. 2002. Cell-cycle control of the establishment of mating-type silencing in S. cerevisiae. Genes Dev 16:2935–2945. doi:10.1101/gad.764102

Lazarus AG, Holmes SG. 2011. A Cis-acting tRNA gene imposes the cell cycle progression requirement for establishing silencing at the HMR locus in yeast. Genetics 187:425–439. doi:10.1534/genetics.110.124099

Lee S, Oh S, Jeong K, Jo H, Choi Y, Seo HD, Kim M, Choe J, Kwon CS, Lee D. 2018. Dot1 regulates nucleosome dynamics by its inherent histone chaperone activity in yeast. Nat Commun 9. doi:10.1038/s41467-017-02759-8

Li YC, Cheng TH, Gartenberg MR. 2001. Establishment of transcriptional silencing in the absence of DNA replication. Science 291:650–3. doi:10.1126/science.291.5504.650

Lindstrom DL, Gottschling DE. 2009. The mother enrichment program: A genetic system for facile replicative life span analysis in Saccharomyces cerevisiae. Genetics 183:413–422. doi:10.1534/genetics.109.106229

Loo S, Rine J. 1994. Silencers and domains of generalized repression. Science 264:1768–1771. doi:10.1126/science.8209257

Martino F, Kueng S, Robinson P, Tsai-pflugfelder M, Leeuwen F Van, Ziegler M, Cubizolles F, Cockell MM, Rhodes D, Gasser SM. 2009. Reconstitution of Yeast Silent Chromatin : Multiple Contact Sites and O - AADPR Binding Load SIR Complexes onto Nucleosomes In Vitro. Mol Cell 33:323–334. doi:10.1016/j.molcel.2009.01.009

McIsaac RS, Silverman SJ, McClean MN, Gibney PA, Macinskas J, Hickman MJ, Petti AA, Botstein D. 2011. Fast-acting and nearly gratuitous induction of gene expression and protein depletion in Saccharomyces cerevisiae. Mol Biol Cell 22:4447–4459. doi:10.1091/mbc.E11-05-0466

Meijsing SH, Ehrenhofer-Murray AE. 2001. The silencing complex SAS-I links histone acetylation to the assembly of repressed chromatin by CAF-I and Asf1 in Saccharomyces cerevisiae. Genes Dev 15:3169–3182. doi:10.1101/gad.929001

Miller A, Yang B, Foster T, Kirchmaier AL. 2008. Proliferating cell nuclear antigen and ASF1 modulate silent chromatin in Saccharomyces cerevisiae via lysine 56 on histone H3. Genetics 179:793–809. doi:10.1534/genetics.107.084525

Miller AM, Nasmyth KA. 1984. Role of DNA replication in the repression of silent mating type loci in yeast. Nature 312:247–251. doi:10.1038/312247a0

Mueller F, Senecal A, Tantale K, Marie-Nelly H, Ly N, Collin O, Basyuk E, Bertrand E, Darzacq X, Zimmer C. 2013. FISH-quant: Automatic counting of transcripts in 3D FISH images. Nat Methods 10:277–278. doi:10.1038/nmeth.2406

Mueller JE, Canze M, Bryk M. 2006. The requirements for COMPASS and Paf1 in transcriptional silencing and methylation of histone H3 in Saccharomyces cerevisiae. Genetics 173:557–567. doi:10.1534/genetics.106.055400

Ng HH, Feng Q, Wang H, Erdjument-Bromage H, Tempst P, Zhang Y, Struhl K. 2002a. Lysine methylation within the globular domain of histone H3 by Dot1 is important for telomeric silencing and Sir protein association. Genes Dev 16:1518–1527. doi:10.1101/gad.1001502

Ng HH, Xu RM, Zhang Y, Struhl K. 2002b. Ubiquitination of histone H2B by Rad6 is required for efficient Dot1-mediated methylation of histone H3 lysine 79. J Biol Chem 277:34655–34657. doi:10.1074/jbc.C200433200

Osborne EA, Dudoit S, Rine J. 2009. The establishment of gene silencing at single-cell resolution. Nat Genet 41:800–806. doi:10.1038/ng.402

Park EC, Szostak JW. 1990. Point mutations in the yeast histone H4 gene prevent silencing of the silent mating type locus HML. Mol Cell Biol 10:4932–4934. doi:10.1128/MCB.10.9.4932.Updated

Picard D. 1994. Regulation of protein function through expression of chimaeric proteins. Curr Opin Biotechnol 5:511–515. doi:10.1016/0958-1669(94)90066-3

Pillus L, Rine J. 1989. Epigenetic inheritance of transcriptional states in S. cerevisiae. Cell 59:637–647. doi:10.1016/0092-8674(89)90009-3

Reiter C, Heise F, Chung H-R, Ehrenhofer-Murray AE. 2015. A link between Sas2-mediated H4 K16 acetylation, chromatin assembly in S-phase by CAF-I and Asf1, and nucleosome assembly by Spt6 during transcription. FEMS Yeast Res 15:fov073. doi:10.1093/femsyr/fov073

Ren J, Wang CL, Sternglanz R. 2010. Promoter strength influences the S phase requirement for establishment of silencing at the Saccharomyces cerevisiae silent mating type loci. Genetics 186:551–560. doi:10.1534/genetics.110.120592

Rine J, Herskowitz I. 1987. Four Genes Responsible for a Position Effect on Expression from HML and HMR in Saccharomyces Cerevisiae. Genetics 116:9–22.

Rufiange A, Jacques PÉ, Bhat W, Robert F, Nourani A. 2007. Genome-Wide Replication-Independent Histone H3 Exchange Occurs Predominantly at Promoters and Implicates H3 K56 Acetylation and Asf1. Mol Cell 27:393–405. doi:10.1016/j.molcel.2007.07.011

Rusche LN, Kirchmaier AL, Rine J. 2003. The establishment, inheritance, and function of silenced chromatin in Saccharomyces cerevisiae. Annu Rev Biochem 72:481–516. doi:10.1146/annurev.biochem.72.121801.161547

Rusché LN, Kirchmaier AL, Rine J. 2002. Ordered Nucleation and Spreading of Silenced Chromatin in Saccharomyces cerevisiae. Mol Biol Cell 13:2207–2222. doi:10.1091/mbc.E02

Santos-Rosa H, Bannister AJ, Dehe PM, Géli V, Kouzarides T. 2004. Methylation of H3 lysine 4 at euchromatin promotes Sir3p association with heterochromatin. J Biol Chem 279:47506–47512. doi:10.1074/jbc.M407949200

Schindelin J, Arganda-Carreras I, Frise E, Kaynig V, Longair M, Pietzsch T, Preibisch S, Rueden C, Saalfeld S, Schmid B, Tinevez JY, White DJ, Hartenstein V, Eliceiri K, Tomancak P, Cardona A. 2012. Fiji: An open-source platform for biological-image analysis. Nat Methods 9:676–682. doi:10.1038/nmeth.2019

Schlissel G, Rine J. 2019. The nucleosome core particle remembers its position through DNA replication and RNA transcription. Proc Natl Acad Sci 116:20605–20611. doi:10.1073/pnas.1911943116

Schneider J, Bajwa P, Johnson FC, Bhaumik SR, Shilatifard A. 2006. Rtt109 is required for proper H3K56 acetylation: A chromatin mark associated with the elongating RNA polymerase II. J Biol Chem 281:37270–37274. doi:10.1074/jbc.C600265200

Siliciano PG, Tatchell K. 1986. Identification of the DNA sequences controlling the expression of the MAT alpha locus of yeast. Proc Natl Acad Sci U S A 83:2320–4. doi:10.1073/pnas.83.8.2320

Steakley DL, Rine J. 2015. On the Mechanism of Gene Silencing in Saccharomyces cerevisiae. G3 (Bethesda) 5:1751–63. doi:10.1534/g3.115.018515

Storici F, Resnick MA. 2006. The Delitto Perfetto Approach to In Vivo Site-Directed Mutagenesis and Chromosome Rearrangements with Synthetic Oligonucleotides in Yeast. Methods Enzymol 409:329–345. doi:10.1016/S0076-6879(05)09019-1

Suka N, Luo K, Grunstein M. 2002. Sir2p and Sas2p opposingly regulate acetylation of yeast histone H4 lysine16 and spreading of heterochromatin. Nat Genet 32:378–383. doi:10.1038/ng1017

Sun ZW, Allis CD. 2002. Ubiquitination of histone H2B regulates H3 methylation and gene silencing in yeast. Nature 418:104–108. doi:10.1038/nature00883

Thorvaldsdóttir H, Robinson JT, Mesirov JP. 2013. Integrative Genomics Viewer (IGV): High-performance genomics data visualization and exploration. Brief Bioinform 14:178–192. doi:10.1093/bib/bbs017

Thurtle DM, Rine J. 2014. The molecular topography of silenced chromatin in Saccharomyces cerevisiae. Genes Dev 28:245–258. doi:10.1101/gad.230532.113

Van Leeuwen F, Gafken PR, Gottschling DE. 2002. Dot1p modulates silencing in yeast by methylation of the nucleosome core. Cell 109:745–756. doi:10.1016/S0092-8674(02)00759-6

van Welsem T, Korthout T, Ekkebus R, Morais D, Molenaar TM, van Harten K, Poramba-Liyanage DW, Sun SM, Lenstra TL, Srivas R, Ideker T, Holstege FCP, van Attikum H, El Oualid F, Ovaa H, Stulemeijer IJE, Vlaming H, van Leeuwen F. 2018. Dot1 promotes H2B ubiquitination by a methyltransferase-independent mechanism. Nucleic Acids Res 46:11251–11261. doi:10.1093/nar/gky801

Wickham H. 2016. ggplot2, Media. New York, NY: Springer New York. doi:10.1007/978-0-387-98141-3

Xu EY, Zawadzki KA, Broach JR. 2006. Single-Cell Observations Reveal Intermediate Transcriptional Silencing States. Mol Cell 23:219–229. doi:10.1016/j.molcel.2006.05.035

Yang B, Britton J, Kirchmaier AL. 2008. Insights into the Impact of Histone Acetylation and Methylation on Sir Protein Recruitment, Spreading, and Silencing in Saccharomyces cerevisiae. J Mol Biol 381:826–844. doi:10.1016/j.jmb.2008.06.059

Young TJ, Kirchmaier AL. 2012. Cell cycle regulation of silent chromatin formation. Biochim Biophys Acta - Gene Regul Mech 1819:303–312. doi:10.1016/j.bbagrm.2011.10.006

